# Cochlear wave dispersion reveals partition micromechanics

**DOI:** 10.64898/2026.11.11.765988

**Authors:** Gabriel Alberts, Sunil Puria

## Abstract

Cochlear traveling waves are a fundamental aspect of mammalian hearing, enabling frequency selectivity. However, incorporating micromechanical interactions between the cells, membranes, and fluids that comprise the cochlear partition presents a challenge when accounting for this selectivity and other macro-scale response properties. The traveling wave can be characterized by its dispersion, which is the relationship between its wavenumber and frequency. One objective was to investigate how organ of Corti structural components influence wave dispersion. To achieve this, we adapted a conventional three-dimensional tapered cochlear box model by incrementally adding morphological characteristics that constitute the partition. Initially a flexible plate, the partition transformed to incorporate a tectorial membrane and inner spiral sulcus. Our secondary objective was to predict the dispersion for each variation using a reduced-length model (slice model). We observed that stiffness and the optimal frequency increased across variations, while wave dispersion remained unchanged relative to the optimal frequency. We also discovered that eigenfrequency analysis, using Floquet boundary conditions, accurately predicted the dispersion for each variation and revealed the existence of other propagating waves including a tectorial membrane transverse mode. These results underscore the significance of analyzing cochlear wave propagation from capturing macromechanical response properties thereby facilitating investigations of micromechanical interactions in reduced-length models.

## I. INTRODUCTION

Theoretical studies of cochlear mechanics typically simplify local tissue interactions to capture global cochlear responses. The cochlear partition, replete with the cells of the organ of Corti between the tectorial and basilar membranes, often gets reduced to a long, thin plate in models than span the full length of the cochlea. While these frameworks have been successful over decades in describing characteristics of the mammalian cochlea, such as its frequency selectivity, they often fail to examine the role of the cochlea’s intricate cytoarchitecture in producing or modifying such characteristics. The cochlear traveling wave underpins the organ’s frequency selectivity and can be described by its dispersion, which relates periodicity in space and time—wavenumber and frequency, respectively. Applying incremental modifications to the cochlear partition to include additional cell types, membranes, and fluid spaces is one way to explore the influence those structural components have on the wave’s propagation. However, creating and solving models that span the length of the cochlea becomes ever more challenging with the addition of each morphological detail that is not tracked in analytical models. Recently, slice finite-element models have been proposed as a means to tackle such a problem (Maftoon et al., 2018; Tubelli et al., 2022; Wang and Puria, 2024; Alberts and Puria, 2024; Kraut and Puria, 2024; Wang and Puria, 2025). Slice models consist of a small cross-section at a specified location of the cochlea with a known best frequency corresponding to that location in the full-length cochlea. This study sought to investigate (a) the impact of incremental additions in cochlear partition morphology on properties like the wavenumber-frequency relationship and (b) how one might predict this relationship from a slice model.

The “box” model, whereby the cochlea is approximated as two equal-area fluid channels separated by a flexible plate, is a well described analytical tool for understanding the cochlea (Peterson and Bogert, 1950). Such models have been used to study a wide range of phenomena including frequency tuning (Steele and Taber, 1979; De Boer, 1981; Neely, 1985; Watts, 2000; Xia et al., 2018), the origins of amplification (Yoon et al., 2011; Wang et al., 2016), dynamic range (Sisto et al., 2021), and the impacts of cochlear implants (Elliott et al., 2011) across numerous species including mouse, guinea pig, chinchilla, cat, and human (Ni et al., 2017). Solving such models requires choosing parameters for the resonant plate like its stiffness and mass.^1^ Studies generally consider stiffness as being derived from the frequency-place map, consistent with the physiological distribution of resonant frequencies from base to apex replicated in a plate-bending-in-fluid model of the cochlea (reviewed in Puria and Steele, 2008). Calculations of mass and its role have been varied with early work taking the fluid mass of the entire scala media (Peterson and Bogert, 1950) and later work suggesting there is no significant mass attached to the basilar membrane (Puria and Steele, 2008). More recent investigations used values that accounted for the thickness of the organ of Corti (Altoè and Shera, 2020; Elliott et al., 2011), although the total value needed to replicate physiological responses changed with the dimensionality of fluid coupling (Steele and Taber, 1979; Elliott et al., 2011).

Traveling waves are a natural outcome of box models since the waves are generated from two properties interacting and the models possess them both: a stiffness gradient and a form of longitudinal coupling (von Bekesy, 1956). While longitudinal coupling exists in many forms (Mammano and Nobili, 1993; Meaud and Grosh, 2010; Yoon et al., 2011; Motallebzadeh et al., 2018; Elliott et al., 2022), we will focus on the dimensionality of fluid coupling for this study as it has been one of the most thoroughly explored forms, yet its effects remain debated. Briefly, 1D fluid coupling produces a pressure difference across the partition that gives rise to a traveling wave whose wavelength decreases with increasing frequency at a single longitudinal place. Allowing the pressure to vary transversely in the fluid chambers (perpendicular to the partition) produces what has been called a “2D short-wave correction.” In this case, pressure and velocity decay with distance from the partition. When the partition width is a small fraction of the scala width—as is considered when applying 3D fluid coupling—the decay from the partition also applies in the width direction (parallel to the partition). Several studies argued that 3D coupling was key to matching physiological data (Steele and Taber, 1979; Kolston, 2000; Lim and Steele, 2002) while others presented lower dimensional solutions for the same problem (Zweig, 2015). A recent theory suggested that although each dimensionality formulation can match experimental results, the pressure and partition admittance required to do so varies with dimensionality (Altoè and Shera, 2020). However, no distinction was made between 2D and 3D solutions. It is currently unclear what degree of coupling is required to capture experimental data and how this may vary from the cochlear base to apex.

Aside from the problem of dimensionality, the field lacks an understanding of how cytoarchitectural components of the organ of Corti shape traveling wave dispersion. While cochlear slice models offer a testbed for such investigations and several versions have been developed, they either did not incorporate a wavenumber-frequency relationship (Ni et al., 2016; Zagadou et al., 2020) or adjusted material parameters to fit data obtained experimentally (Cai and Chadwick, 2003; Cai et al., 2004). Furthermore, it has been shown that multichamber cochlear models produce multiple modes of wave propagation beyond the classical traveling wave which could influence amplification and inner hair cell stimulation (Steele, 1999; Steele and Puria, 2005). Although some modal studies reverted to the box model geometry (Watts, 2000; Elliott et al., 2013), one recent study involving a 3D slice model reported a mode which deformed the organ of Corti and produced longitudinal fluid flow within it (Zagadou et al., 2020), potentially a mechanism for outer hair cell motility to amplify OoC motion. Despite the promising results, there remains no description of how the constituent components of the partition affect the dispersion relation. Such a description would help clarify the mechanical roles of these structures and build intuition for the future design of more detailed cochlear slice models.

Overall, our goal is to bridge the gap between the macro and micromechanical understanding of cochlear mechanics. We have approached this in two ways: 1) by adapting traditional, full-length box models to include additional levels of morphological detail, and 2) by developing corresponding slice models. We proceed by comparing the full-length and slice models for each morphological variation with increasing complexity. Specifically, we first explore the response of the tapered full-length model with no structures attached to the basilar membrane and how its dispersion compares to each dispersion calculation using the corresponding slice model. We then add stepwise to that model to include an arch in the basilar membrane, a pseudo-organ of Corti, and a tectorial membrane with an inner spiral sulcus and subtectorial fluid spaces. Each study investigates the effects of a given partition modification on the wavenumber-frequency relationship as well as on stiffness, passive cochlear tuning, the frequency-place map, and displacement and pressure gain. In addition, we tested whether the wavenumber-frequency relationship from each variation could be predicted using only information from the corresponding cochlear slice model. We investigated the effects of stiffness and mass with one-, two-, and three-dimensional longitudinal fluid coupling based on asymptotic assumptions to determine which combination best reflected the dispersion relation obtained across model variations.

## II. FINITE-ELEMENT METHODS

A full-length, finite-element tapered box model of the gerbil cochlea was developed which contained numerous morphological features of the cochlear partition: an arch-beam representation of the basilar membrane (BM) (Kapuria et al., 2017; Xia et al., 2018), a solid mass above to represent a pseudo-organ of Corti (OoC), and a tectorial membrane (TM) complete with a fluid inner spiral sulcus (ISS) (Fig. 1). The model also included periodic fluid subtectorial spaces (STS) which acted as a channel between the ISS and the larger scala media/vestibuli and periodic solid structures that connected the OoC and TM representing the coupling from outer hair cell bundles. While cochlear morphology has been studied thoroughly, incorporating details like individual hair cells and hair bundles would make the computation ever more prohibitive in a full-length model. This is in fact a primary motivation for the use of cochlear slice models (Wang and Puria, 2025). Since one of the goals of this investigation was to test techniques that predict the dispersion relation of cochlear slice models, we sought to develop a model with enough detail in geometry, physics, and materials to probe such techniques. To that end, we considered four morphological features to be the most relevant as each has been found or hypothesized to play an important role in cochlear wave dispersion: scala height, BM stiffness, OoC mass, and a fluid chamber within the cochlear partition. Taking scala height, for example, the model was tapered from base to apex, which has been shown to be important for characterizing the cochlear input impedance (Puria and Allen, 1991; Shera and Zweig, 1991). While the BM is typically considered a flexible plate due to its collagen fibers, the gerbil exhibits a unique arch (Edge et al., 1998; Kapuria et al., 2017; Xia et al., 2018), which was also considered in this model. The model was considered sufficient once the authors felt each of the four features was addressed.

**FIG. 1.**
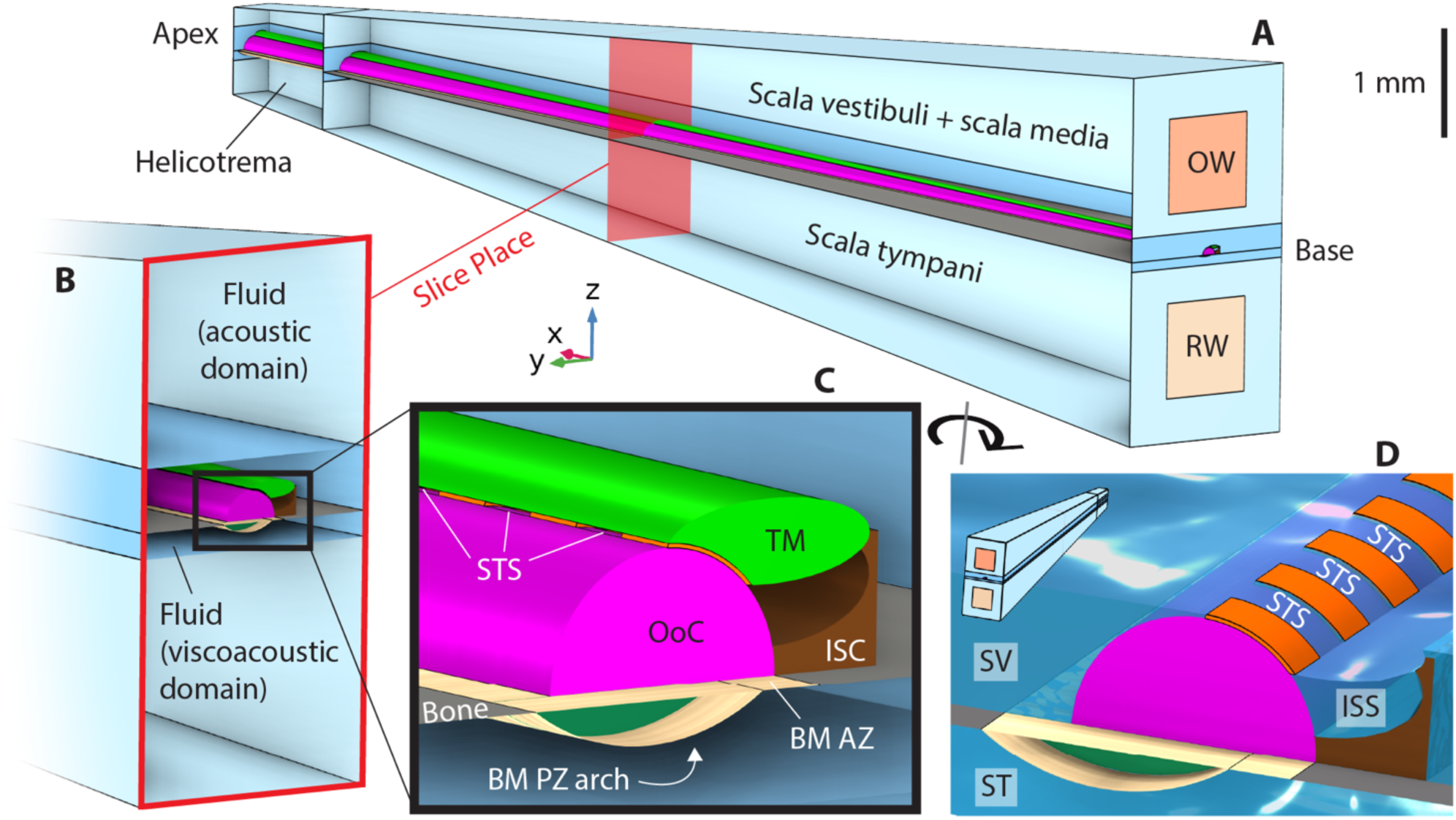
(Color online). Full-length finite-element tapered box model morphology. A) Full-length model based on the gerbil cochlea with base to the right and apex to the left. Longitudinal, radial, and transverse directions correspond to the x, y, and z axes, respectively. Scala vestibuli was combined with scala media into one fluid space which joined scala tympani through an opening in the helicotrema (no gray bone next to BM PZ arch). A velocity source was specified at the oval window (OW) and zero pressure condition at the round window (RW). The slice place was 7.575 mm from the base corresponding to a best frequency of 2500 Hz. B) View of the model cut at the slice place showing the subdivision of the fluid into far-field acoustic and near-field viscoacoustic domains for computation. C) Zoomed-in view of the cochlear partition with a basilar membrane (BM) split between an arcuate zone (AZ) and arched pectinate zone (PZ), pseudo-organ of Corti (OoC), and tectorial membrane (TM) with longitudinally periodic fluid subtectorial spaces (STS). The BM, which was filled with ground substance in the volume formed by the radial and arch fiber layers, was attached to bone on either end. Solid material is shown deeper than bone and BM to highlight the thin BM arch and overlap of the OoC and BM arcuate zone. The TM was connected to the OoC by longitudinally periodic structures (orange) interleaved with the STS that approximately represented the mechanical coupling from outer hair cell bundles. The TM was also connected to a structure representing inner sulcus cells (ISC) and limbus. D) Rotated partition view with TM removed to show coupled fluid spaces. Scala vestibuli (SV) and scala tympani (ST) were above and below the partition, respectively. The inner spiral sulcus (ISS) was bounded by the TM (hidden for clarity), ISC, and OoC. The longitudinally spaced STS fluid channels connect the ISS and SV. The inset in the top left shows the full-length model in the new orientation.

We also made a corresponding finite-element cochlear slice model on which we performed an eigenfrequency analysis. The slice model was constructed using the properties at a single longitudinal distance from the base in the full-length model—the same place used for the dispersion calculation in the full-length model. Importantly, we examined the cumulative effects of adding morphological features starting with a flexible plate only and culminating in the model variation with the TM and ISS. Since each variation built on the one prior, geometric and material properties for a given morphological feature, for example the basilar membrane arch, are consistent once introduced with no additional parameter tuning. Therefore, the model type presented in Fig. 1 and described below refers to the final variation with the TM, ISS, and all other morphological features. A section following these descriptions details how each variation was constructed from the one prior.

### A. Full-length Box Model

A full-length finite-element model of the gerbil cochlea was developed in COMSOL Multiphysics version 6.0 which included numerous morphological features of the cochlear partition including a tectorial membrane (TM) and inner spiral sulcus (ISS). Geometrical and material properties of the model including width (y-direction) of the BM, OoC and TM area, and BM Young’s modulus were based on previous literature. A complete accounting of parameters is presented in Table I.

**TABLE I.**
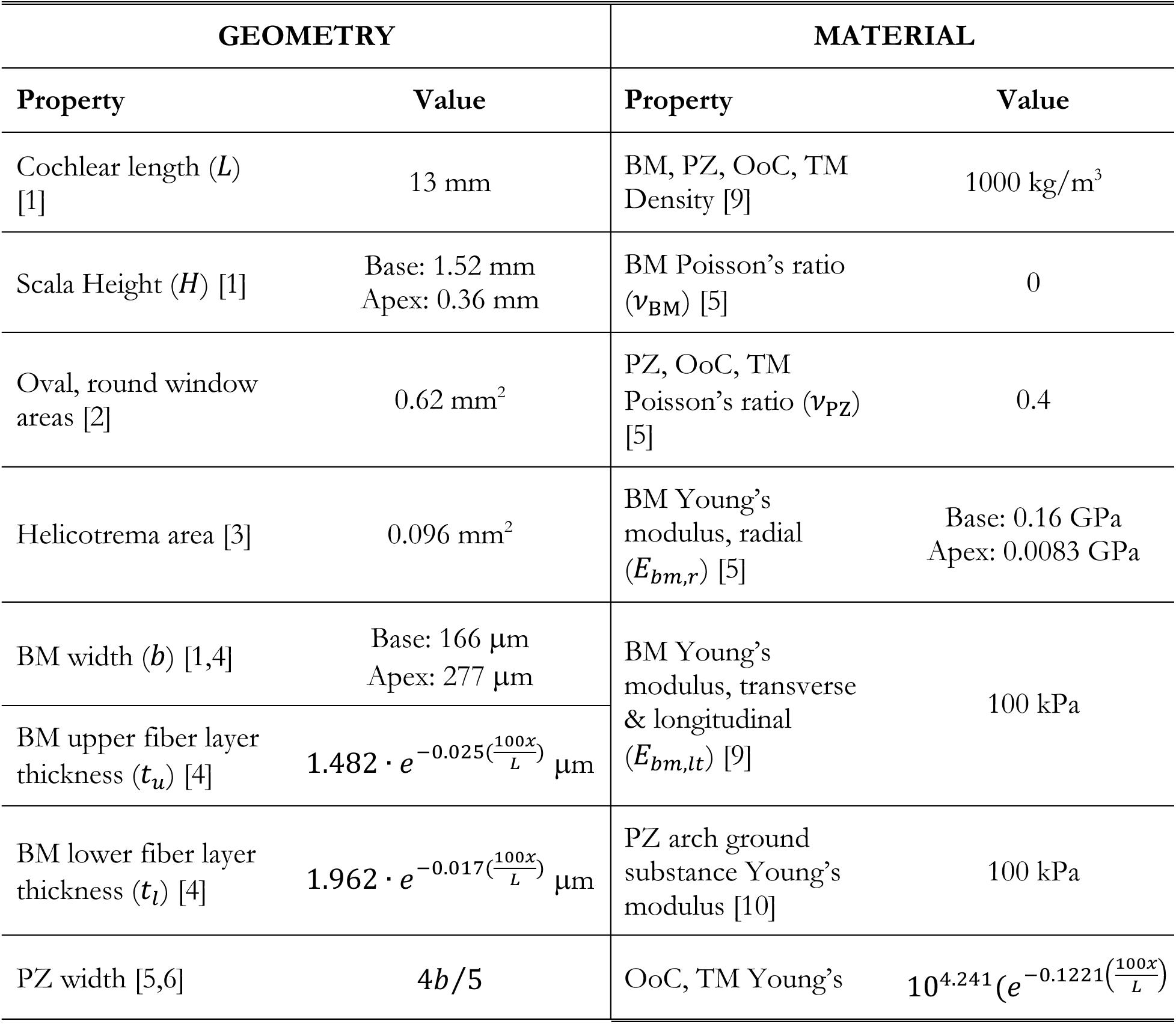
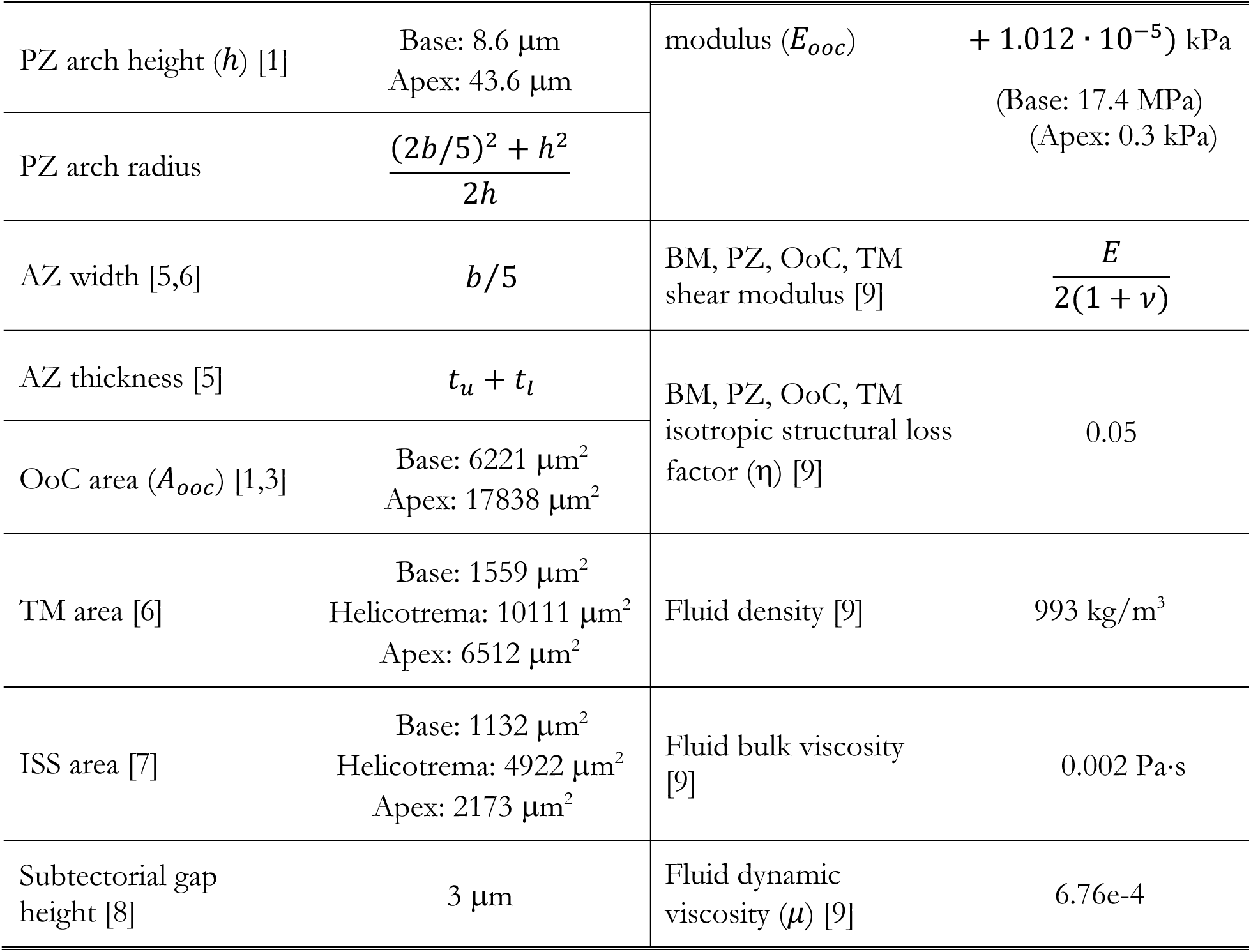
Full-length box model geometric and material properties. *x* is distance from the cochlear base. ‘Base’ refers to *x* = 0 and ‘Apex’ refers to *x* = *L*. Values were linearly interpolated between locations except for the TM and ISS areas for which the asymmetric faces were lofted between locations resulting in slightly smaller areas compared to linear interpolation. All references to Plassmann et al. come from measurements of the gerbil species *Meriones unguiculatus*. The shear modulus was calculated using the Young’s modulus from the respective structure and orientation when applicable. Fluid material properties come from liquid water in the built-in COMSOL Materials library. References: [1] (Plassmann et al., 1987), [2] (Dong and Olson, 2006), [3] (Mountain et al., 2003), [4] (Schweitzer et al., 1996). [5] (Kapuria et al., 2017), [6] (Edge et al., 1998), [7] (Cho et al., 2022), [8] (Fettiplace, 2017), [9] (Motallebzadeh et al., 2018), [10] (Naidu and Mountain, 2007).

| GEOMETRY |  | MATERIAL |  |
|---|---|---|---|
| **Property** | **Value** | **Property** | **Value** |
| Cochlear length (*L*) [1] | 13 mm | BM, PZ, OoC, TM Density [9] | 1000 kg/m<sup>3</sup> |
| Scala Height (*H*) [1] | Base: 1.52 mm<br>Apex: 0.36 mm | BM Poisson’s ratio (ν<sub>BM</sub>) [5] | 0 |
| Oval, round window areas [2] | 0.62 mm<sup>2</sup> | PZ, OoC, TM Poisson’s ratio (ν<sub>PZ</sub>) [5] | 0.4 |
| Helicotrema area [3] | 0.096 mm<sup>2</sup> | BM Young’s modulus, radial (*E*<sub>*bm,r*</sub>) [5] | Base: 0.16 GPa<br>Apex: 0.0083 GPa |
| BM width (*b*) [1,4] | Base: 166 μm<br>Apex: 277 μm | BM Young’s modulus, transverse & longitudinal (*E*<sub>*bm,lt*</sub>) [9] | 100 kPa |
| BM upper fiber layer thickness (*t*<sub>*u*</sub>) [4] | $1.482 \cdot e^{-0.025(\frac{100x}{L})}$ μm |  |  |
| BM lower fiber layer thickness (*t*<sub>*l*</sub>) [4] | $1.962 \cdot e^{-0.017(\frac{100x}{L})}$ μm | PZ arch ground substance Young’s modulus [10] | 100 kPa |
| PZ width [5,6] | 4*b*/5 | OoC, TM Young’s modulus (*E*<sub>*ooc*</sub>) | $10^{4.241}(e^{-0.1221\left(\frac{100x}{L}\right)} + 1.012 \cdot 10^{-5})$ kPa<br>(Base: 17.4 MPa)<br>(Apex: 0.3 kPa) |
| PZ arch height (*h*) [1] | Base: 8.6 μm<br>Apex: 43.6 μm |  |  |
| PZ arch radius | $\frac{(2b/5)^2 + h^2}{2h}$ |  |  |
| AZ width [5,6] | *b*/5 | BM, PZ, OoC, TM shear modulus [9] | $\frac{E}{2(1 + \nu)}$ |
| AZ thickness [5] | *t*<sub>*u*</sub> + *t*<sub>*l*</sub> | BM, PZ, OoC, TM isotropic structural loss factor (η) [9] | 0.05 |
| OoC area (*A*<sub>*ooc*</sub>) [1,3] | Base: 6221 μm<sup>2</sup><br>Apex: 17838 μm<sup>2</sup> |  |  |
| TM area [6] | Base: 1559 μm<sup>2</sup><br>Helicotrema: 10111 μm<sup>2</sup><br>Apex: 6512 μm<sup>2</sup> | Fluid density [9] | 993 kg/m<sup>3</sup> |
| ISS area [7] | Base: 1132 μm<sup>2</sup><br>Helicotrema: 4922 μm<sup>2</sup><br>Apex: 2173 μm<sup>2</sup> | Fluid bulk viscosity [9] | 0.002 Pa·s |
| Subtectorial gap height [8] | 3 μm | Fluid dynamic viscosity (*μ*) [9] | 6.76e-4 |

#### 1. Geometry

The cochlear length was taken as the average of the inner and outer BM lengths (Plassmann et al., 1987) to approximate the uncoiled cochlea (Fig. 1A). Reisner’s membrane was considered acoustically transparent, so scala vestibuli and scala media were combined into one fluid domain (Motallebzadeh et al., 2018; Zweig et al., 1976) equal in area with scala tympani, both with square cross sections for simplicity. To capture general trends in geometric variations along the cochlear length, scala area as well as OoC area, BM width, and pectinate zone arch height were extrapolated from linear fits to data from the middle 80% of the cochlea (Plassmann et al., 1987; Schweitzer et al., 1996). The flexible BM was centered radially between the scalae walls with fixed bone on either side (Fig. 1B). The only exception was in the helicotrema (apical 10% of the cochlear length) where the bone adjacent to the BM pectinate zone was removed to connect the fluid scalae. The oval window (stapes footplate) and round window were equal area square cutouts positioned at the cochlear base in the center of each scalae for geometric simplification and computational efficiency.

Given the dispute over the effective mass of the cochlear partition, we prioritized accounting for the solid mass of the tissues and fluid mass of the enclosed fluid spaces by making our best effort to preserve PZ height in addition to OoC, TM, and ISS cross-sectional area (Fig. 1C). This allowed us to compare theoretical values of fluid mass against the mass of the solid tissues and enclosed fluids for a given morphological variation.

Since the challenges associated with developing full length models grow with the inclusion of each morphological detail, we homogenized the cells of the OoC with the tunnel of Corti and outer tunnel fluid into a single solid mass in the shape of a half cylinder, similar to Tubelli et al. (2025). The cylinder radius was a function of distance from the base, so the cross-sectional area mirrored that of measured morphological data. Likewise, the thousands of outer hair cell hair bundles firmly connecting the OoC and TM in a typical mammalian cochlea were approximated by longitudinally periodic solid sections attached to the OoC and TM which extended radially from the ISS to the scala (orange sections in Fig. 1D). These were interleaved with periodic fluid channels that formed subtectorial spaces which connected the ISS and scala fluids.

The ISS was bounded by the OoC, TM, and another structure that represented the inner sulcus cells and limbus. This structure was included to provide a fixed, solid connection for the “modiolar” region of the TM away from the OoC radially and to separate the ISS and scala fluid spaces so their only connection was through each STS.

#### 2. Materials

The density for all soft tissue was 1000 kg/m^3^—near that of water at gerbil body temperature (38 °C). The basilar membrane upper and lower fiber layers were transversely orthotropic with a radial Young’s modulus equal to the modulus for collagen, taken to be 1 GPa, times the volume fraction of collagen, which decreased from 0.16 at the base to 0.0083 at the apex (Kapuria et al., 2017)^1^. The BM Young’s modulus in the transverse and longitudinal directions was over three orders of magnitude lower at 100 kPa, equal to that of the PZ ground substance (Naidu and Mountain, 2007).

Since the pseudo-OoC combined multiple cell types and fluids into one structure, the structure required material properties which accounted for such a combination. Hence, the OoC was made isotropic with a Young’s modulus based on tissue properties (Akhtar et al., 2011) and further refined so the frequency-place map was near a half-octave below the in-vivo map from Müller (1996), roughly as expected for a postmortem, passive cochlea.^2^ For simplicity, the TM and inner sulcus cell support structure were also isotropic and given the same Young’s modulus as the OoC, varying from base to apex. At our analysis location in the middle turn of the gerbil cochlea (2.5 kHz best place), the TM Young’s modulus was 14 kPa, similar to previously reported values (Freeman et al., 2003; Steele and Puria, 2005).

Importantly, none of the parameters changed between studies of varying cochlear partition morphologies. By not retuning parameters, we were able to observe the effects of stepwise additions of morphological features like a change in partition stiffness, tuning sharpness, gain, or dispersion. For example, this allowed us to determine whether the OoC contributes any substantial stiffness to the cochlear partition or whether the stiffness arises purely from the BM. Keeping parameters consistent across variations helped clarify the mechanical roles of each structure and build our intuition for the future design of more intricate cochlear slice models.

#### 3. Physics

The model fluid was split into two domains: a near-field thermoviscous acoustic domain and far-field pressure acoustics domain to reduce computation time (Motallebzadeh et al., 2018). Viscous losses are concentrated at the partition (Wang et al., 2016), so the viscoacoustic domain included the entire ISS and all STS as well as a region surrounding the partition. The height of this region was made greater than the boundary layer thickness at the lowest frequency tested—29 μm at 250 Hz. While thermal effects were captured, they were not of interest for the studies presented in this manuscript. An inviscid fluid approximation was used for the scala vestibuli and scala tympani volumes outside of the region farther from the partition.

Tissues were also split between two domains: shell structures for the BM and bone, and solid structures for the PZ, OoC, and TM. The BM fiber layers were modeled as 2D shells (surfaces) to reduce computational cost as they exhibit significant bending stiffness and are thin compared to their width and length along the cochlea. Fixed bone on either side of the BM was also made using shells to avoid narrow fluid regions around the solid-shell connections which were not of interest. The other structures could not be approximated in a similar fashion and were modeled using the complete 3D formulation.

The divisions across fluids and structures allowed us to accurately capture the motion of each with boundary-layer losses while retaining computational efficiency. Equations describing the various physics are detailed in the appendix.

#### 4. Boundary Conditions and Multiphysics Coupling

Scalae walls were rigid aside from the oval and round windows. The oval window was prescribed a velocity of 0.01 mm/s for all frequencies across all morphological variations. The round window was prescribed 0 Pa as expected for an open middle ear cavity (Wang and Puria, 2025). The centered, flexible partition was clamped on either side to a fixed boundary as bone has been shown to move much less than the BM (Johnstone et al., 1970; Rhode, 1971). The structure that borders the ISS and connects to the “modiolar” edge of the TM (radially opposite the OoC) was fixed on the surfaces facing the scala, representing a rigid limbus. The entire partition was fixed at the basal and apical ends of the model.

Coupling between the inviscid and viscous domains was prescribed by continuity in normal acceleration and stress. Between the fluid-structure and structure-structure interfaces, continuity was enforced in velocity and displacement. Coupling equations are provided in the appendix.

#### 5. Mesh and Computational Configurations

The mesh was semi-automatically constructed across domains to ensure adequate resolution in narrow regions while maintaining computational feasibility. The viscous boundary layer thickness largely determined the minimum mesh density required and was defined as 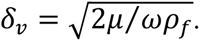 For water at gerbil body temperature and atmospheric pressure, this was approximately 10 μm at 3.25 kHz and larger at lower frequencies. The fluid mesh surrounding the measurement place was fine enough to contain at least three nodes within the boundary layer for frequencies up to 3.25 kHz. This did not resolve the layer for higher frequencies and thus underestimated viscous losses there. However, those frequencies were presented only for completeness of the place map and were not used for the dispersion studies, which allowed us to save development and computation time. The STS, where most viscous loss has been predicted to occur (Allen, 1980; Prodanovic et al., 2015), was the narrowest region in the model and had seven nodes (three quadratic elements) across each dimension (height, width, and length) of each STS fluid channel. The final mesh contained 1,014,901 elements and 7,857,409 degrees of freedom. The model converged for all frequencies tested.

Computations were made on a Dell Precision Tower 7910 computer running Ubuntu 18.04.5 LTS on a Linux kernel with two 20-core Intel(R) Xeon(R) CPUs (E5-2698 v4) and 512 GB of RAM. The model was solved in the frequency domain with simulations taking 62 minutes per frequency.

### B. Floquet Slice Model

Floquet-Bloch theory separates geometric periodicity from wave periodicity. While originally applied to study electron transport in crystals (Bloch, 1929), the theory has been used to evaluate wave propagation in fields such as acoustic metamaterials (Fleury et al., 2016) and seismology (Colombi et al., 2016) and for applications including nondestructive testing (García and Fernández-Álvarez, 2015; Hakoda et al., 2026). In practice, a ‘unit cell’ is evaluated which represents one of an infinite spatial array of unit cells. The cell size is independent of the various wavelengths considered during evaluation. This saves valuable computational resources and development time when analyzing cells with intricate details. The Floquet slice model was our unit cell (Fig. 2). Despite being 3D, Floquet boundary conditions were applied only to the longitudinal faces of the slice separated by Δ*x*. This restricted wave propagation and morphological repetition to the longitudinal (x) direction, effectively creating an infinitely long cochlea. Importantly, discontinuities between longitudinal faces could be avoided while still incorporating longitudinal variations of structures, materials, and physics *within* the slice model. The Floquet slice model and its relation to the cochlear traveling wave are presented in Figure 2.

**FIG. 2.**
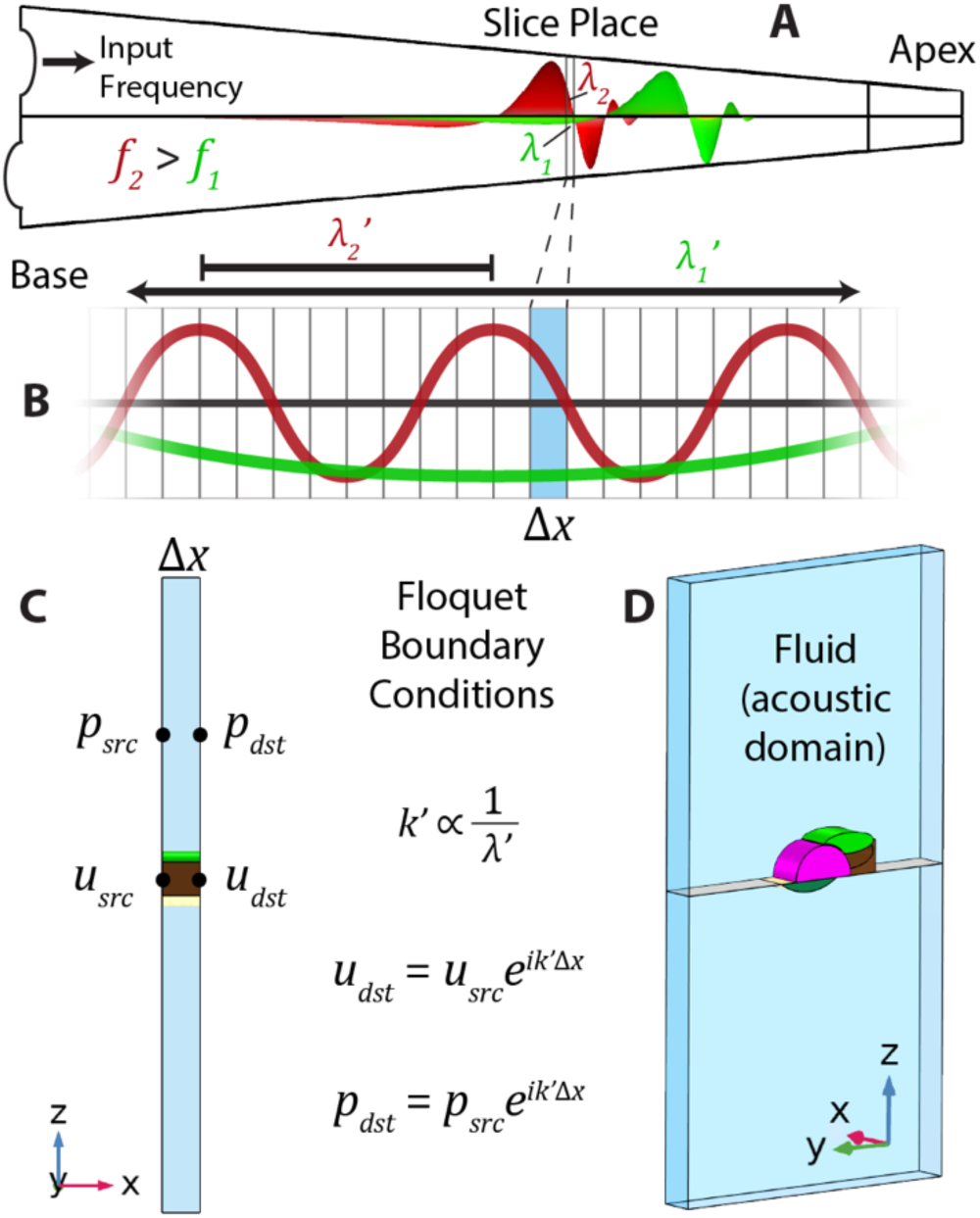
(Color online). Cochlear traveling wave and the Floquet slice model. A) Full-length model traveling wave profile for two different frequency inputs f_1_ and f_2_ (f_2_ greater than f_1_) showing the longitudinal wavelength shortening with increasing distance from the base. The local wavelength (λ_1_, λ_2_) can be determined from the phase change across a short longitudinal span (two vertical gray lines). B) Floquet boundary conditions simulate an infinite repeating array of slices (vertical rectangles) with an infinite repeating wave whose wavelength is independent of the geometric periodicity of the slice. Shown are two wavelengths λ_1_′, λ_2_′ approximately equal to the local wavelengths λ_1_, λ_2_ in (A) where λ_2_′ is shorter than λ_1_′. C) Longitudinal-transverse (x-z) view of the Floquet slice model with slice thickness Δ*x*. To simulate wave propagation, Floquet boundary conditions define a phase shift in pressure *p* and velocity *u* across the slice thickness (x-direction) from a source face to a destination face that corresponds to a given wavelength that is inversely proportional to wavenumber *k* ′. D) View of the Floquet slice model in a similar orientation to Fig. 1B. The slice contains the same morphological features as the full-length model. All fluid domains were inviscid as viscosity was excluded for eigenfrequency analysis.

The Floquet slice version of the full-length model variation described in II.A. had properties taken from a single longitudinal cross section of the full-length model—the same place analyzed for the wavenumber-frequency relationship. Material and geometric properties were made uniform across the slice thickness such that the only longitudinal variation was the alternating sections of solid pseudo-hair bundles and fluid STS. Since these each extended 50 μm longitudinally and repeated one after the other, the slice was made 100 μm thick—the smallest possible length that captured all morphological features.

Identical physics from the full-length model were used except for the replacement of viscous fluid with inviscid fluid. Viscosity was expected to have a small effect on the real component of the wavenumber (Steele and Taber, 1979; Puria and Steele, 2008), which was negligible when considering a slice model (Wang and Puria, 2025). As will be discussed, including viscosity was not necessary to predict the dispersion of the full-length models.^3^ Hence, it was not included for computational and post-processing efficiency. Boundary conditions and physics couplings were also identical to the full-length model where possible (see appendix for new coupling equations). The longitudinal truncation meant there was no stapes velocity, round window pressure, or partition fixation at the basal or apical ends, and instead only Floquet boundary conditions applied to the longitudinal faces. Contrary to the full-length model frequency domain studies, our Floquet slice model was evaluated with an eigenfrequency study, which is used to solve for the resonant frequencies of a system with no forcing function. Therefore, unlike Wang and Puria (2025), the model had no stimulation condition. The eigenfrequency analysis is described further in III.D. The final mesh for the slice—130 times shorter than the full-length model—had 17,418 elements and 74,909 degrees of freedom. Each simulation took 22 seconds per wavenumber.

### C. Models for Other Cochlear Partition Morphological Variations

We examined the cumulative effects of adding morphological features to the cochlear partition starting with only a flexible plate and culminating in the model variation with the TM and ISS described thus far. Since the other variations each had fewer morphological features, they were built by deconstructing the TM/ISS variation (Fig. 3). The pseudo-OoC model was generated by replacing the TM, solid TM-OoC connections, and inner sulcus cell structure with viscous fluid and modifying physics couplings to reflect the new fluid-structure boundaries. The arch BM model was similarly constructed from the pseudo-OoC model simply by replacing the OoC with viscous fluid and updating physics couplings to reflect the change. In the case of the flat BM model, the ground substance of the arch BM was deleted so the upper and lower fiber layers combined into a flat partition as in the arcuate zone of the arch BM model. As we were interested in exploring the effects of presenting a given morphological feature, no parameters were adjusted with any partition modification. Solver configurations were unchanged aside from the frequencies solved for. The slice place was consistent between variations to eliminate effects from longitudinal differences in parameters, so the best frequency (BF) for that place was determined by iteration in each variation. Frequencies solved were relative to the BF. The Floquet slice model for the other variations was generated in a similar fashion to the methods detailed above for the TM variation of the full-length box model.

**FIG. 3.**
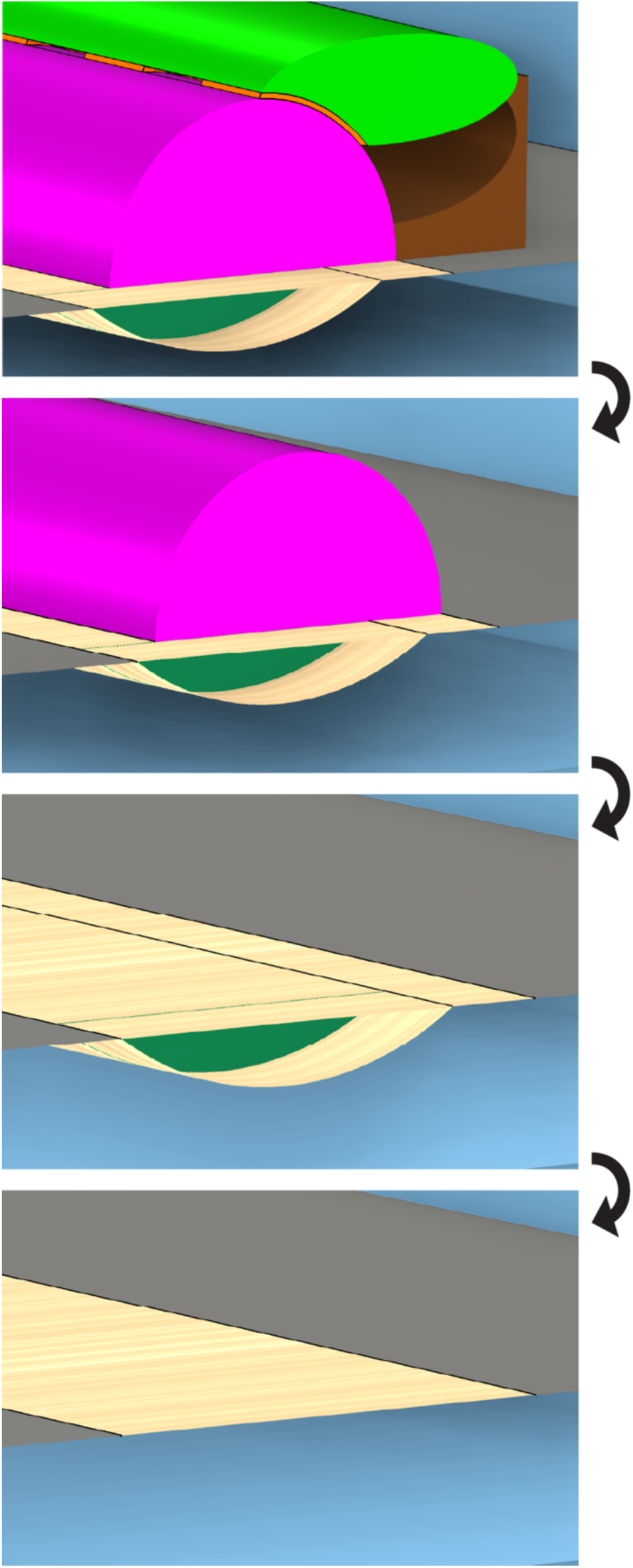
(Color online). Morphological variations formed by deconstructing the cochlear partition of the full-length model from Fig. 1. Four partition variations were studied including the baseline ‘TM/ISS’ model previously described (top to bottom): TM/ISS, pseudo-OoC, arch BM, and flat BM.

## III. DISPERSION CALCULATION METHODS

Three formulations of the dispersion relation for a given slice model were investigated to compare against the dispersion obtained from the corresponding full-length model. The first two apply the 2D Wentzel-Kramers-Brillouin (WKB) approximation to probe the effects of stiffness, mass, and viscosity. The first followed the methods described in (Puria and Steele, 2008) for inviscid and viscous fluid using only the partition stiffness as obtained numerically from a static load on the slice model partition. The second considered the complex impedance of the partition and made use of a fixed-point algorithm to arrive at a solution after iteration as described in (Frost, 2024) and used in (Altoè et al., 2021; Altoè and Charaziak, 2023; Altoè and Shera, 2020; Shera and Altoè, 2023). With damping set to zero (De La Rochefoucauld and Olson, 2007), comparing the first two approaches provided a direct evaluation of the role of mass in passive cochlear tuning. Lastly, we obtained dispersion curves by performing an eigenfrequency study on the Floquet slice model complete with all geometrical and material parameters and structural and fluid domains as described in II.B. Since three-dimensional fluid coupling effects influence the cochlear base where the fraction of the BM width to scala width is small (Altoè, 2026) and our place of interest was the middle turn of the gerbil cochlea, our analysis focused on one- and two-dimensional analytical descriptions of fluid coupling. What follows are the steps underlying each dispersion calculation method, starting with the baseline calculation for the full-length models.

### A. Dispersion from Full-length Model

The dispersion relation of a full-length box model was taken from the slope of the BM transverse velocity phase with respect to longitudinal place:

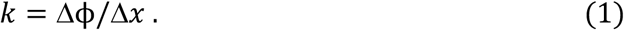

The phase was measured at the arcuate-pectinate junction in each model with an arch BM although the slope did not change when measured at other radial locations along the BM (not shown). In the flat BM model, the slope was measured from the center of the partition. The slope was evaluated +/- 50 μm around the place of interest for each frequency for which the model was solved. This calculation was similar to those made for experimental data (De La Rochefoucauld and Olson, 2007).

### B. Dispersion from Slice Model, Method 1: Volume Stiffness

Puria and Steele (2008) argued that for mechanical purposes there is no significant mass attached to the BM. We used the 2D representation they derived to calculate the dispersion of a cochlear slice model from its volume stiffness. Volume stiffness was defined as the pressure difference across the partition (transverse, scala vestibuli to scala tympani) divided by the cross-sectional transverse area displacement of the partition

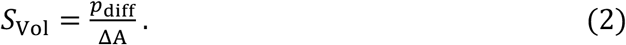

This is equivalent to the area-normalized stiffness (pressure over vertical BM displacement in specific acoustic impedance units) divided by the BM width. We obtained quantitative values for pressure and displacement using a static pressure study of our finite-element slice models (Supplementary Fig. S3). We applied a 1 Pa pressure load to the ST-facing BM then integrated the transverse displacement along that BM surface and divided by the slice thickness to obtain a BM cross-sectional area displacement. This allowed for any longitudinal variations in geometry, materials, or physics to be accounted for. While the only such change in the studies presented here was the variation in STS, the procedure could be used, for example, for more detailed slice models with individual hair cells.

The dispersion relation came from the transmission line wave equation, which, for a single frequency and variables changing in time as *e^jωt^*, reduces to

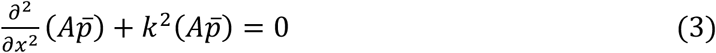

with

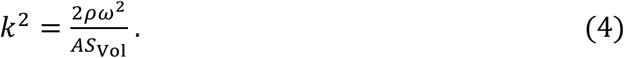

Here, *A* is the cross-sectional area for a single scala (assuming symmetric scalae), *p̄* is the average pressure for a single scala, and *k* is the local wavenumber. The factor of two accounts for pressure from each scala. The positive and negative roots of *k* represent forward and backward traveling waves, but only forward going waves are explored here.

This 1D approximation assumes equal pressure throughout the scala and is valid in the long-wave region of the cochlea. However, it breaks down near the best-frequency, short-wave region because the pressure along the BM is much greater than the average scala pressure (Olson, 1998). The short-wave pressure gain can be expressed following (Duifhuis, 1988) as

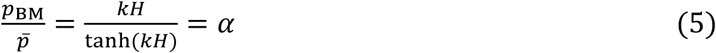

which approaches one for small *kH* and goes as *kH* for large values of *kH*. While Eq. (3) still holds, the wavenumber is modified such that

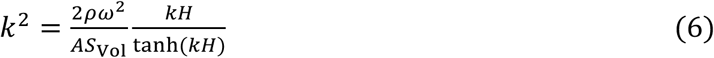

Taking the scala cross-sectional area to be square so *A* = *H*^2^, Eq. (6) can be approximated to a long- and short-wave region based on the limiting behavior of the hyperbolic tangent function with

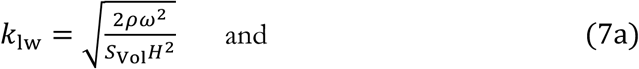

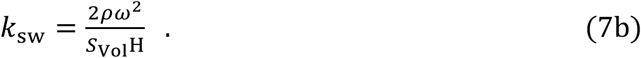

Equations (7a) and (7b) are solutions for the case of inviscid fluid. To incorporate fluid viscosity *μ*, a parameter β was defined 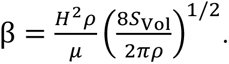 For large β, the wavenumber becomes

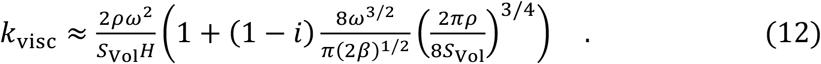

Compared to the short-wave wavenumber, the effect of viscosity was to increase the real component and produce a negative imaginary component, although we were only concerned with the real component which dictated the wavelength.

To quantitatively evaluate these variables, we took *H* to be the height of the scala at the place of interest, density *ρ* and dynamic viscosity μ based on the material parameters ( 1000 kg/m^3^ and 7e-4 Pa·s, respectively), and *S*_Vol_ as calculated above. The transition frequency between the long- and short-wave regions occurred at 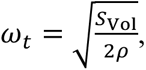 so the long-wave solution was appropriate for frequencies below this value and the short-wave solution for frequencies above. The three wavenumbers (*k*_lw_, *k*_sw_, and *k*_visc_) are presented for each morphological variation.

### C. Dispersion from Slice Model, Method 2: Complex Impedance

Shera and Altoe (2023) presented a formulation for the wavenumber of an active cochlea as a modification of the passive, complex wavenumber. Since the present work considers only the passive case, we will use the approach as a foundation that could be extended in future studies to address active mechanics. The passive wavenumber in the long-wave region is defined as

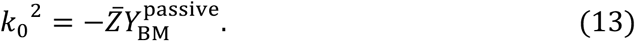

with *Z̄* the fluid impedance for square scalae *Z̄* = 2*iωρ*/*H*^2^ in units of acoustic impedance over length. Additionally, 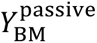 is the passive admittance of the BM modeled as a simple harmonic oscillator which, in units of length over specific acoustic impedance, was

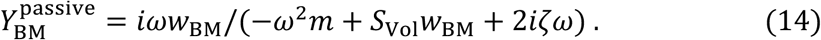

where *m* was the area-normalized OoC mass, *w*_BM_ was the BM width, and *ζ* the damping factor which was set to zero for these studies as damping’s influence on passive mechanics is negligible. Various masses were considered by multiplying the density (same for fluid and solids) by the volume of a particular structure or set of structures and dividing by the slice width. For a stiffness-dominated admittance, the wavenumber is equivalent to Eq. (7a) as expected.

Short-wave pressure gain was accounted for using the gain factor *α* such that for passive mechanics *k*^2^ = *αk*_0_^2^. This resembled Eq. (6) now with mass effects included. Notably, since *k* and *α* were mutually dependent, the equation was transcendental and the wavenumber required a numerical solution. We followed a fixed-point algorithm as described by (Frost, 2024) and used in (Shera and Altoè, 2023) to iterate between solutions for *k* and *α*

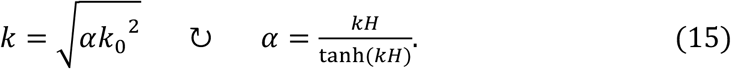

until the wavenumber varied by less than 1%—our chosen criteria and the same as (Altoè and Shera, 2020)—which typically required less than eight iterations. Results were presented for combinations of solid and fluid masses with the structure denoted in the subscript of the line name (e.g. m_OoC_ uses the mass of the pseudo-organ of Corti). Computations for methods one and two were performed in MATLAB (R2023a, The MathWorks, Natick, MA).

### D. Dispersion from Slice Model, Method 3: Eigenfrequency

Eigenfrequency studies are used to solve for the resonant frequencies of a system with no forcing function. Associated eigenmodes provide a representation of the response shape with an undefined magnitude. Acoustic waveguides, for example, are commonly described by their propagating modes. These modes display varying shapes, are associated with a particular propagating wavenumber, and are frequency dependent. The Floquet boundary conditions create a virtual waveguide with a fixed propagating wavelength from which the propagating frequencies can be computed. Therefore, we employed an eigenfrequency study using the Floquet slice model equipped with all structures and fluid spaces to evaluate the dispersion relations native to the slice. In general, the homogeneous, linearized, governing differential equations were solved for a single wavenumber according to

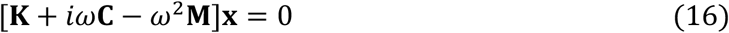

where **K**, **C**, and **M** were the global stiffness, damping, and mass matrices, respectively, and **x** the eigenvectors. While eigenfrequencies are traditionally solved for by setting det(**K** + *iω***C** − *ω*^2^**M**) = 0 (Van Der Heijden, 2014), more computationally efficient strategies were used given the much larger finite-element matrices (see supplemental material for solver specifications). Since different wavenumbers produced different matrices, the wavenumber-frequency relation was obtained by prescribing a series of Floquet wavenumbers and solving an eigenfrequency study of the slice model at each wavenumber. The dispersion relation was then the combined results of the eigenfrequency studies across wavenumbers. Structural damping in the system contributed to the imaginary component of the resulting eigenfrequencies, but our investigations only considered the real components, which represented angular frequencies.

## IV. RESULTS

We will first explore the classic response of the tapered full-length model with no structures attached to the BM and how this dispersion compares to each dispersion calculation using the corresponding slice model. We will subsequently build the finite-element models (full-length and corresponding slice) towards that shown in Fig. 1 complete with a BM arch, pseudo-OoC, and TM. Through each variation, we will investigate the effects of a given partition modification and how to account for the effects in the dispersion calculation. We examined the dispersion at the 7.575 mm place roughly 60% from the base as we sought to study the gerbil middle turn and avoid effects unique to the very base and apex. The location corresponded to the 2.5 kHz place in the TM model variation shown in Fig. 1 and was kept constant across the other model variations.

### A. Flat BM

We begin with the box model at its most simple geometric construction where the centered, flexible partition consisted of only a flat BM with no solid structures attached. Spatial and frequency responses of the flat BM full-length model along with dispersion calculations using the full-length and slice models are presented in Fig. 4. While the full-length model nominally included viscosity, the dispersion for an equivalent inviscid full-length model was also shown to tease apart effects explicitly due to viscosity. All other results including the velocity transfer function, displacement and pressure gains, and place map refer strictly to the full-length model with viscosity. Similar models have been described in the literature (Steele and Taber, 1979; De Boer and Van Bienema, 1982; Neely, 1985; Elliott et al., 2011; Ni et al., 2017), and our results are consistent with their findings.

**FIG. 4.**
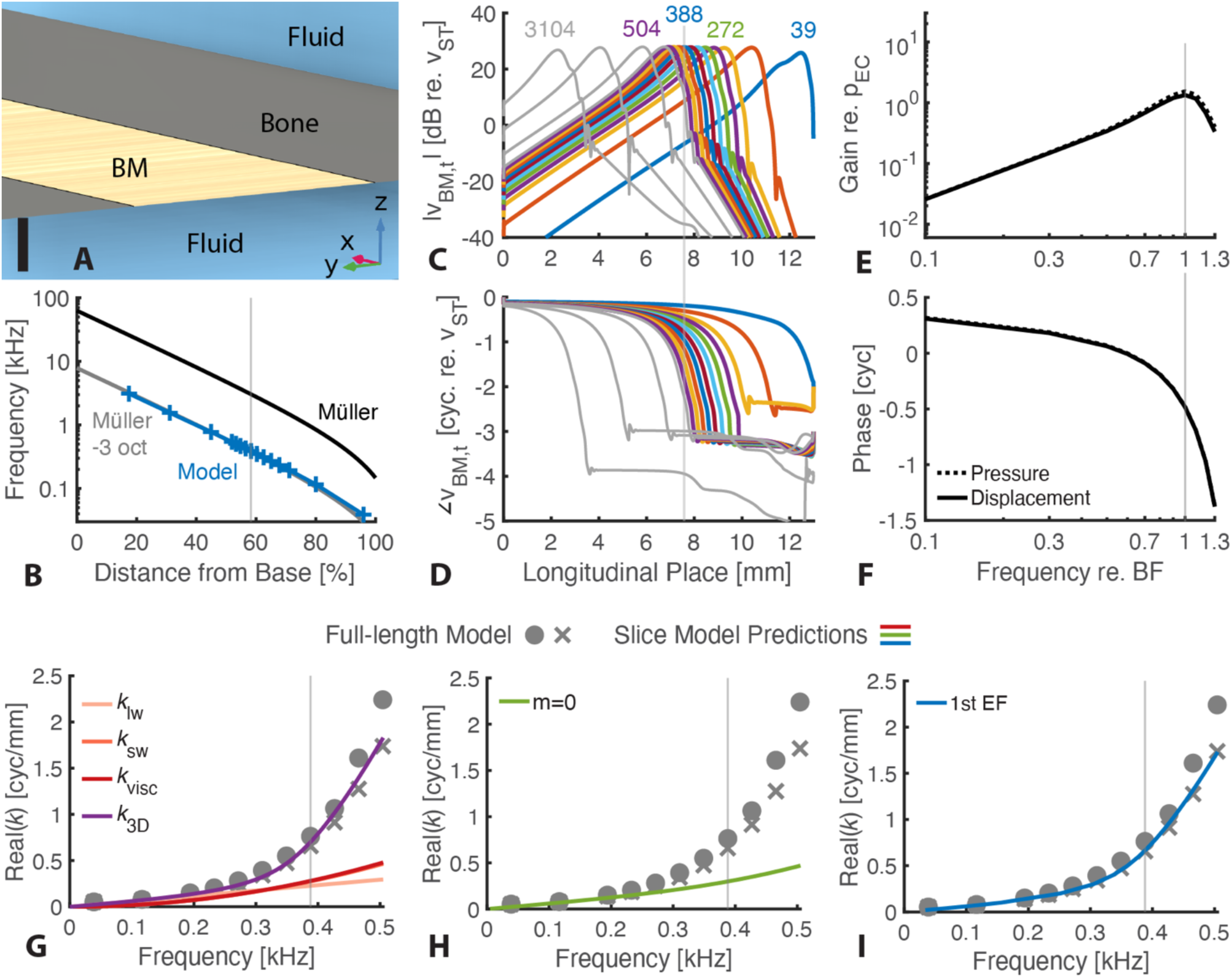
(Color online). Flat BM full-length model response and dispersion calculations. A) Model partition geometry at the 7.575 mm place showing the flexible BM in the center, bone on either side radially (y-direction), and fluid above and below. Fluid immediately surrounding the partition was removed for viewing the underlying structures. Scale bar 50 *μ*m. Panels B-F correspond to results from the full-length model with viscosity. An inviscid model was constructed solely for wavenumber analysis of this partition morphology (panels G-I). B) Place-frequency map with the vertical axis frequency in kHz and horizontal axis longitudinal place in percent distance from the base. The vertical line indicates the longitudinal location analyzed for subsequent plots E-I. The Müller map (1996) was shown (black line) with *CF*(*pos*) = 0.398 · 710^(100–*pos*)·0.022^ − 0.631; where *pos* is percent distance from the base and *CF* is in kHz. Alongside is the map shifted three octaves down (gray line) for comparison to the model (plus markers) since, for this flat BM partition, the best frequency (BF) near 60% from the base was 388 Hz. C,D) Transverse velocity magnitude (C) and phase (D) versus longitudinal place. Partition velocity, measured at the center of the BM, is plotted with respect to stapes velocity. Responses are shown in color for frequencies used in the dispersion calculation: 0.1, 0.3, and (in steps of 0.1) 0.5 to 1.3 times the best frequency (BF) of the place of interest around 7.5 mm from the base. Responses are shown in gray for higher frequencies used to fill the frequency-place map: 1.4, 2, 4, and 8 times the BF. The vertical line is centered about the place of interest and is 100 μm thick—the longitudinal span over which the model’s phase slope was computed for the dispersion calculation shown as the circles in panels G-I. E,F) Frequency response magnitude and phase of local pressure difference (dotted lines) and BM displacement (solid lines) relative to ear canal pressure. Values were obtained using the input stapes velocity and a high pass filter to approximate the middle ear transfer function based on Ravicz et al. (2008). The local pressure difference was measured 1 μm above and below the center of the partition into SV and ST, respectively. Horizontal axis: frequency relative to best frequency (BF). Vertical axis: displacement gain in μm/Pa and pressure gain (dimensionless). A positive pressure difference was defined to produce a negative displacement, so displacement phase was shifted 0.5 cycle to offset this definition and better compare displacement and pressure. Displacement and pressure responses overlapped across frequencies. G-I) Real part of the wavenumber (cycles per millimeter) vs frequency (kHz) for three slice calculation methods (colored lines) and two full-length models (viscous (circles) and inviscid (x-marks)). The wavenumber from the full-length models was calculated as the negative slope of the velocity phase about the place of interest. The BF of the viscous model was indicated by the vertical line (388 Hz) while the BF of the inviscid model was about a quarter octave higher (463 Hz). The wavenumber including viscosity was less than 1.3 times higher than the inviscid wavenumber for all frequencies shown. G) Wavenumbers calculated using method 1 (volume stiffness) including the long wave, short wave, and viscous solutions (light to dark, respectively). A stiffness-only calculation with 3D fluid coupling is included for reference and is discussed further in Supplemental Material section IV.E. H) Wavenumber from method 2 (complex impedance iteration method) using no mass since no tissue surrounds the BM. I) Wavenumber from method 3 (eigenfrequency). Only one eigenfrequency (EF) was within the frequency range of interest for all wavenumbers.

The model’s tonotopy resulted in a place map (Fig. 4B) that covered roughly 7 octaves in frequency^4^. This span of frequencies was similar to the *in vivo* place map (Müller, 1996). However, the absolute frequencies were shifted three octaves down. While most models select parameters to match experimental responses like neural tuning curves (Neely, 1985), BM transfer functions (Altoè and Shera, 2020), or frequency-place maps (Elliott et al., 2011), the present model was based entirely on anatomical parameters. Our results suggest that the BM width, thickness, and fiber density alone are insufficient to produce the higher frequency responses seen in the gerbil cochlea.

The normalized velocity magnitude (Fig. 4C) and phase (Fig. 4D) were similar to those of previous box models. The magnitude increased exponentially around 6 dB/mm in the basal region before peaking near 25 dB. In this region, the model exhibited the classic traveling wave phase accumulation up to around four cycles. At the base, the phase was around -0.2 cycles due to a combination of the cochlear input impedance phase and model orientation^5^. Similar results were found in other cochlear models (Tubelli et al., 2025). Apical to the peak, the magnitude decayed rapidly and the phase reached a plateau suggestive of influence from the fast compression wave (see supplemental material section III.A for details).

The passive displacement and local pressure gain relative to ear canal sound pressure level exhibited a bandpass frequency response (Fig. 4E,F). Using a high-pass middle ear admittance with a 2 kHz cutoff frequency (see Supplemental Material section IV.C.), the frequency response profile was similar to the stapes-normalized BM velocity response in Fig. 4C,D^6^. With no frequency dependence between the model’s pressure and displacement, the cochlear response was stiffness-dominated, as expected through the BF (Altoè et al., 2022). The displacement gain peaked near 1000 nm/Pa while the pressure gain peaked near 1 indicating the local pressure difference across the partition at BF was equal to the ear canal sound pressure level. The low frequency phase was near 0.25 cycles due to the phase of the middle ear admittance and accumulated ¾ cycles of phase by the BF.

The wavenumber-frequency relationships obtained from the three slice calculation methods described in Section III are presented in Fig. 4G-I, each compared against full-length model dispersion. The wavenumber was evaluated through 1.3 times the BF to capture the short-wave response prior any phase plateau or ripples apical to the peak (Altoè and Charaziak, 2023; De La Rochefoucauld and Olson, 2007). The BF of the viscous model occurred around ¾ cycles/mm or a wavelength of 1.3 mm, similar to experimental data from the gerbil middle turn (Zosuls et al., 2021).

Figure 4G shows these results compared to the 2D WKB solution considering solely the partition stiffness. The model approximated the long-wave solution well at the lowest frequencies (*k*_lw_). However, even the inviscid full-length model wavenumber increased beyond the short-wave approximation (*k*_sw_) starting around an octave below the BF. Viscosity was expected to slightly increase the wavenumber with a larger effect at higher frequencies as evidenced by the inviscid versus viscous full-length models. While the predicted wavenumber including viscosity (*k*_visc_) increased compared to the short-wave approximation, it remained well below the dispersion curves of the inviscid and viscous full-length models. While the model included no solid mass (Fig. 4A), several studies have described how a layer of fluid rides with the partition and can be interpreted as an added mass in a 3D inviscid model (Steele and Taber, 1979; De Boer, 1981; Neely, 1985). This is relevant for our model as the partition occupied a fraction of the scala width. The 3D dispersion (*k*_3D_) overlaps with the dispersion of the inviscid full-length model well. However, this was not true for the other model variations, so the 2D results are discussed (see Supplemental Material section IV.E. for details regarding 3D dispersion). Since the partition included no solid mass, the wavenumber from the complex impedance in Fig. 4H was equivalent to the combined long- and short-wave 2D approximation from Fig. 4G.

Lastly, Fig. 4I showed the full-length model dispersion curves compared to those computed using the eigenfrequency method described in Section III.D. The Floquet slice model (inviscid) produced a single eigenfrequency in the range of interest for all prescribed wavenumbers (solid line), which overlapped directly with the curve for the inviscid full-length model. The partition displacement was that of a clamped plate peaking at the center, as seen in the full-length model (Supplemental Material section IV.A.). The fluid pressure distribution was equal throughout each scala at low wavenumbers and low frequencies and was more concentrated near the partition as the wavenumbers and frequencies increased, also mirroring the response from the full-length model. This remarkable agreement shows that the local wavenumber in a tapered full-length flat BM cochlear model can be well approximated by solving for the eigenfrequencies of a local slice model using Floquet boundary conditions.

### B. Arch BM

We next considered the unique nature of the gerbil basilar membrane, which consists of an arched pectinate zone (PZ) in addition to a flatter arcuate zone (AZ) (Edge et al., 1998; Kapuria et al., 2017; Xia et al., 2018). The full-length model responses and dispersion calculations from the full-length and slice models are presented alongside an illustration of the new morphology in Fig. 5. Since the inviscid Floquet slice model captures the dispersion of the full-length model with viscosity (Fig. 5I), there is no additional inviscid full-length model for comparison. This holds for the current arch BM morphology variation as well as the OoC and TM/ISS variations to come.

**FIG. 5.**
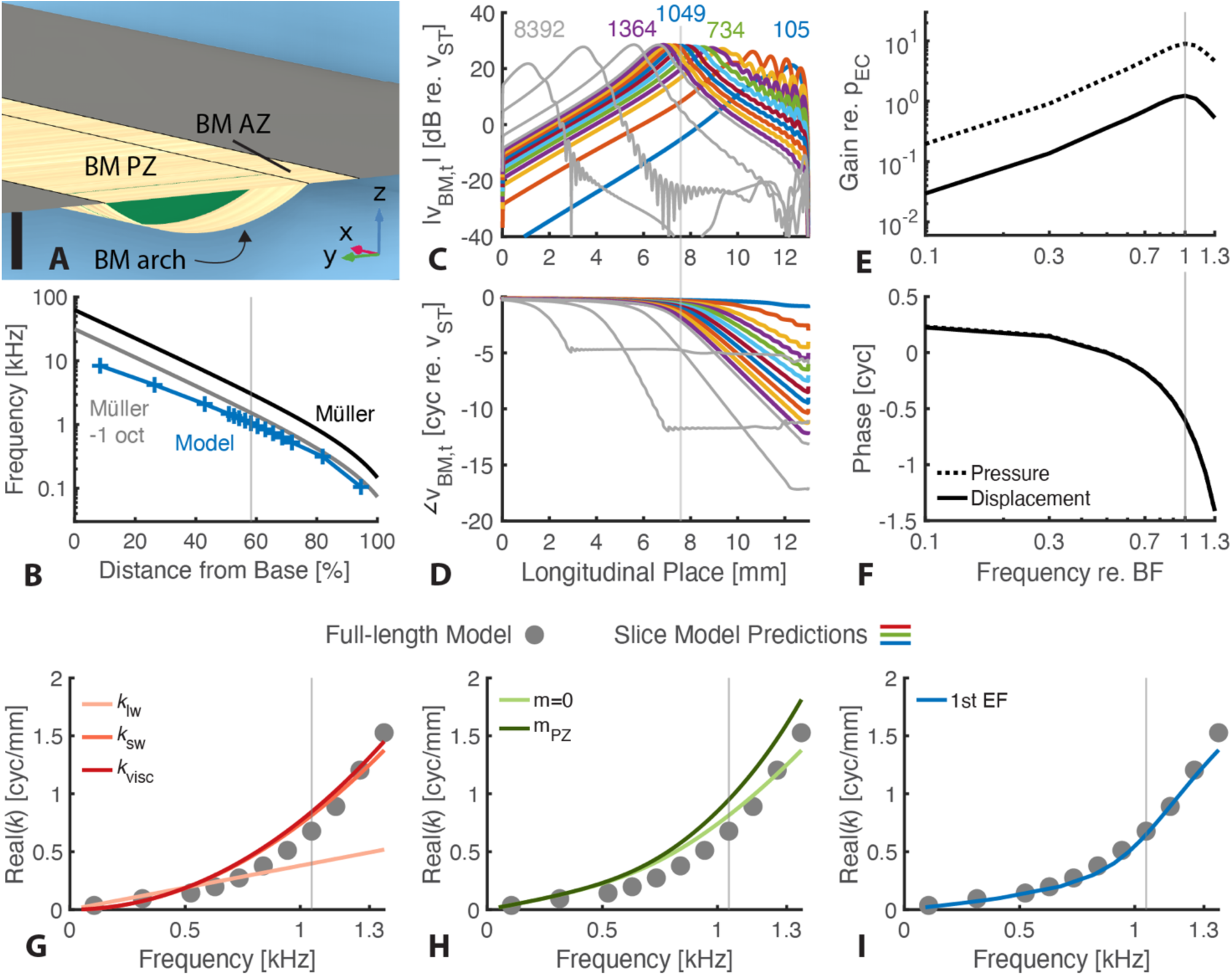
(Color online). Arch BM model response and dispersion calculations. The figure is structured similarly to Fig. 4 and will only be elaborated upon here. A) Model partition morphology at the 7.575 mm place, now showing the flexible BM split into two regions: an arcuate zone (AZ) and arched pectinate zone (PZ) with the PZ comprising 80% of the BM width. The PZ ground substance (green), was shown at a more apical distance than the BM to highlight the thin fiber layers that surround the substance. Scale bar 50 *μ*m. B) Place-frequency map showing the arch BM model (plus markers) compared to the in-vivo Müller map (black line) and the map shifted one octave down (gray line). C, D) Normalized transverse velocity magnitude (C) and phase (D) measured at the junction between the arched PZ and flat AZ. The same frequencies relative to BF were used as the flat BM model but now the BF at the place of interest was 1049 Hz. E,F) BM displacement (solid lines) and local pressure (dotted lines) gain frequency response relative to ear canal pressure. Displacement was measured between the AZ and PZ and pressure was measured 1 μm above and below this junction into SV and ST, respectively. G-I) Wavenumber vs frequency plots against full-length model dispersion (dots). G) Stiffness-only long-wave, short-wave, and viscous wavenumber approximations. H) The complex impedance iteration dispersion showing the case for no mass (light line) and the added mass of the PZ ground substance (dark line). I) Eigenfrequency analysis of the Floquet slice showing the lowest acoustic eigenfrequency (EF).

Compared to the flat BM, the introduction of an arch in the BM fiber layers produced a slightly more flattened place map, which was now mostly within two octaves of the in vivo map and closer to one octave at lower frequencies (Fig. 5B). At our place of interest roughly 60% from the base, the BF was 2.7 times higher relative to the flat BM variation at 1049 Hz. Taking the stiffness of the two variations as calculated from a static pressure load (section III.B and supplementary material section II.C.), the partition stiffness increased by a factor of 2.5. Since tissue density is near that of fluid, a resonance-based place map primarily concerns the change in stiffness across the length of the cochlea. Therefore, partition resonance would increase with square root of stiffness. Here, the expected resonance change from stiffness alone was a factor of 1.6, less than the factor of 2.7 measured BF change.

The addition of the BM arch produced a similar velocity magnitude and phase profile until just past the peak where the magnitude remained higher and the phase continued accumulating (Fig. 5C,D). The phase plateau only appeared for the highest frequencies suggesting that the fast wave only dominated once the velocity magnitude fell 30 dB below that of the stapes. The arch broke the radial symmetry of the flat BM model and skewed the velocity radial profile (velocity along the BM width, y-direction). The velocity now peaked at the junction between the AZ and PZ (shown in supplemental material section IV.A.), which replicates experimental findings (Cooper, 2000; Cooper and Van Der Heijden, 2018).

The displacement and pressure difference gains maintained a similar bandpass shape to the flat BM model relative to BF since the stimulus frequencies were still below the 2 kHz middle ear cutoff frequency (Fig. 5E,F). However, while the displacement gain was quantitatively similar, the pressure gain increased by close to a factor of seven (2.7x increase in frequency multiplied by 2.5x change in stiffness, see supplemental material section IV.C.). While the change in partition stiffness could have resulted in a lower displacement, higher pressure, or combination of the two, for this modification we found the displacement re. ear canal pressure was undisturbed while the local pressure gain increased.

The dispersion curve for the arch BM full-length model was again nonlinear with a wavenumber at BF around ¾ cycles/mm, similar to the flat BM model. The wavenumber calculations, however, revealed a different origin for the response. Figure 5G showed that the 2D approximation based solely on the partition stiffness closely predicted the wavenumber found in the full-length model. This meant the added fluid mass from 3D fluid coupling was no longer necessary and overestimated the wavenumber despite the BM width and scalae dimensions remaining identical to the flat BM variation (supplementary material section IV.E.). Including the mass of the pectinate zone ground substance kept the 2D approximation similar to the full-length model (Fig. 5H).

The eigenfrequency study accurately captured the differences from the flat BM case and again produced a single eigenfrequency in the frequency range of interest for all wavenumbers, which corresponded to the traveling wave mode (Fig. 5I). As in the full-length model, the displacement peaked at the junction between the AZ and PZ and pressure focused around this point at higher frequencies and wavenumbers. The tight overlap between the dispersion of the viscous full-length model and inviscid Floquet slice show the negligible effects of viscosity on the traveling wave wavenumber.

### C. Pseudo-Organ of Corti

The OoC contains numerous cell types of varying sizes, orientations, and material properties in addition to fluid spaces like the tunnel of Corti and outer tunnel that vary along the length of the cochlea. However, developing and computing a full-length model with such detail becomes ever more prohibitive in a full-length model, which motivates the use of cochlear slice models. To minimize computational complexity and capture the area occupied by the tissues and fluid, the OoC was simplified to a solid semicylinder on top of the BM extending into scala vestibuli. Unlike the ground substance mass in the pectinate zone arch, the pseudo-OoC was not surrounded by the stiff BM fiber layers. Frequency responses for this morphological variation are shown in Fig. 6 alongside dispersion curves comparing the full-length model results and slice model predictions.

**FIG. 6.**
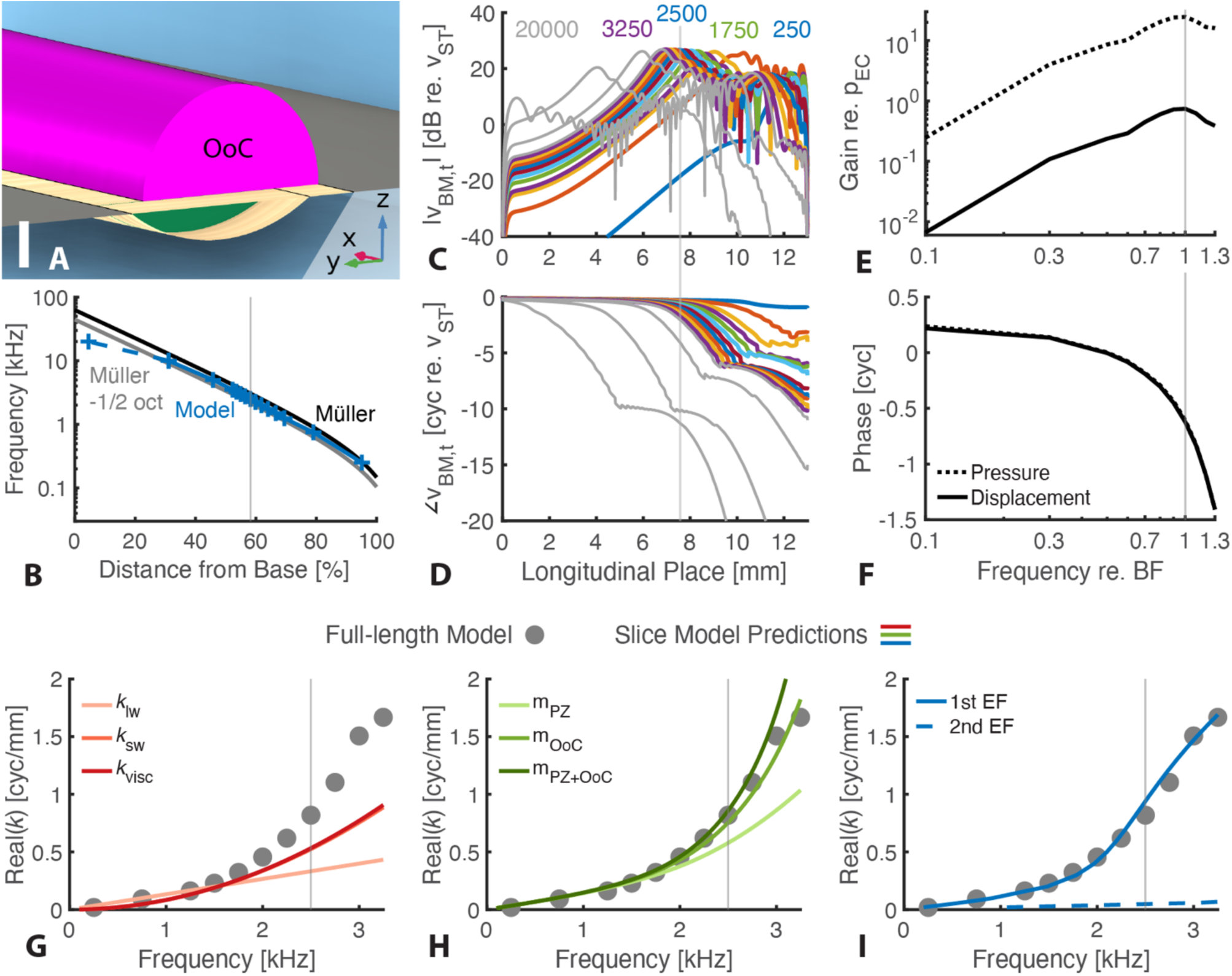
(Color online). OoC model response and dispersion calculations. The figure structurally mimics Fig. 4 and 5 so will only be elaborated upon. A) Partition morphology now includes a semicylindrical OoC in magenta atop the BM intersecting with the AZ-bone interface. The pseudo-OoC cross-sectional area was 3.6 times larger than the PZ area. Scale bar 50 *μ*m. B) Place-frequency map showing the model nearly a half-octave-shifted from the in-vivo Müller map. The BF was now 2500 Hz. The highest frequency was connected via dashed line as it indicates the place of the second highest peak since an apical wave produced a larger response, shown in C. E,F) BM displacement was measured between the arcuate and pectinate zones. ST pressure was measured 1 μm below the junction, and SV pressure was measured 1 μm above the center of the OoC. G-I) Wavenumber vs frequency plots. H) The complex impedance iteration method showed the dispersion curves predicted from including the PZ mass only, OoC mass only, and combined mass (light to dark green). I) The lowest acoustic eigenfrequency (solid line) again fully captured the slow traveling wave wavenumber, but an additional mode in the frequency range of interest was present (dashed line) which corresponded to a compression of the partition.

The place map overall resided roughly a half octave below the in vivo Müller map (Fig. 6B). The isotropic, place-dependent Young’s modulus of the OoC was chosen to restore the range of the in vivo map and the half-octave shift—roughly as expected for a passive cochlea. Compared to the arch BM model, the BF at our place of interest was higher by a factor of 2.3 and was now 2500 Hz. At the ∼60% place, the OoC Young’s modulus was about 30 kPa—slightly more compliant than the ground substance in the BM arch and much more compliant than the BM fibers. However, despite this relatively compliant structural addition, the stiffness (determined by static pressure load on the ST-facing BM) was higher by a factor of 9.6. This change in stiffness predicts a 3.1x increase in BF based on a stiffness-dominated resonance place map, more than the 2.3x change determined from the full-length models. The BF shift could be better accounted for with a 1.8x increase in effective mass in addition to the stiffness change, although this value did not correspond to the mass of the OoC tissue relative to the PZ ground substance since the OoC cross-sectional area was 3.6x larger.

The spatial velocity response showed a slightly steeper growth toward the peak for a given frequency compared to the previous studies while the peak value itself was nearly unchanged (Fig. 6C,D). Ripples in the magnitude and phase responses apical to the maximum were more prominent compared to previous morphological variations but were not considered further as the frequency for dispersion analysis was limited to just above the BF. A second peak apical to the maximum is similarly not considered for these investigations except to explain the highest frequency on the place map (connected by dashed line) where the basal peak (20 kHz marker) exhibited a lower velocity magnitude than the secondary apical peak.

Since the middle ear transfer function had a cutoff frequency of 2 kHz, the displacement and pressure gains showed broader peaks around BF, which was determined by the unnormalized local BM velocity. This meant the displacement and pressure were not boosted as much near BF relative to ear canal pressure as for the previous morphological variations. The resulting displacement gain was roughly three times lower near BF and the pressure gain higher compared to the arch BM case. Unlike from the introduction of a BM arch, the change in partition stiffness now resulted in a lower displacement and higher local pressure gain because of the shift in frequency. Displacement and pressure responses still overlapped, signifying the stiffness-dominated cochlear response.

The dispersion curve for the full-length model including the pseudo-OoC was shown as circles in Fig. 6G-I and was qualitatively similar to the other studies relative to the BF. In this case, the stiffness was appropriately captured as can be seen by the low-frequency, long-wave approximation overlapping with the full-length model results (Fig. 6G). However, the full-length model dispersion deviated from the stiffness-only short-wave approximation at higher frequencies with viscosity providing a meager improvement. For the pseudo-OoC, the dispersion relation was much more accurately captured when incorporating its mass in the 2D approximation (Fig. 6H). The dispersion from the full-length model largely landed between those incorporating only the OoC mass and both the OoC and PZ mass. In the eigenfrequency analysis, the traveling wave mode was again captured across frequencies by the lowest eigenfrequency including the decreasing slope above BF. Additionally, a second mode appeared in the frequency range of interest where the ST and SV-facing boundaries moved out of phase, compressing and expanding the OoC. The in-phase pressure across each scala was reminiscent of the fast wave, although the slope of the curve gave a speed of 50 m/s, far below the 1500 m/s speed of sound propagation in water.

### D. Tectorial Membrane and ISS Fluid Tunnel

So far, we have seen how the wavenumber can be quantitatively similar relative to the BF across variations despite being dominated by fluid mass, stiffness, or solid mass depending on the partition morphology. Our final morphological variation included a TM and ISS fluid space, which allowed us to investigate the effects of the added solid and fluid mass and the effects of adding a fluid tunnel while leaving the remaining structures from the previous studies intact. The normal physiology of the mammalian cochlea includes the TM sitting atop rows of stereociliary bundles of outer hair cells and was greatly simplified in this case. Alternating 50 μm longitudinal sections of solid and fluid below the TM created structural coupling between the OoC and TM and fluid coupling between the ISS and SV via the STS. The TM edge opposite the OoC was fixed, representing the limbus, and an additional solid structure was placed to create the remaining boundary of the ISS. As with the previous studies, model responses and wavenumber predictions are presented (Fig. 7).

**FIG. 7.**
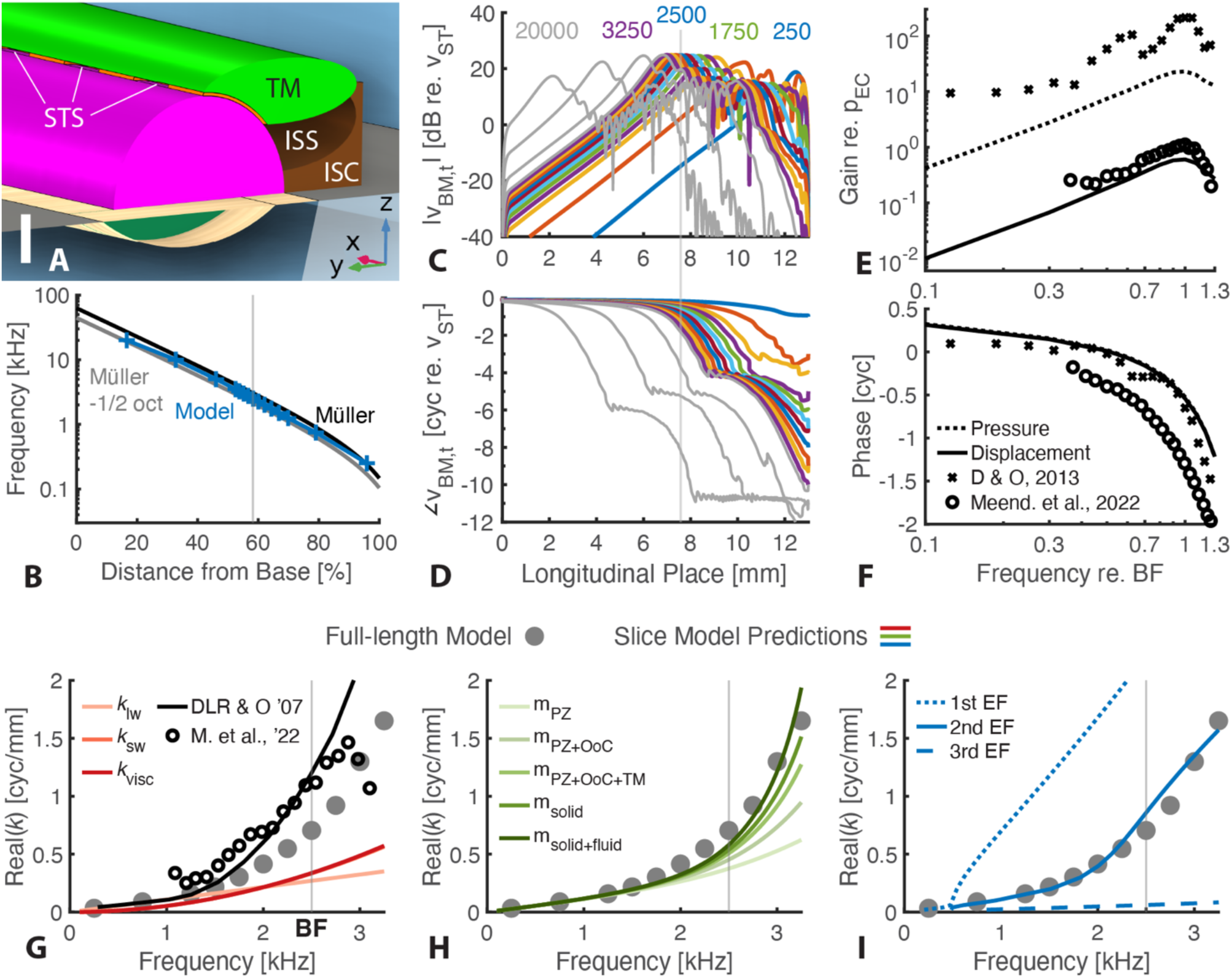
(Color online). TM/ISS Model responses. Deviations from Fig. 6 are discussed. A) Model partition morphology shows the addition of the tectorial membrane (TM, green) and inner sulcus cells (ISC), which form the boundaries of the inner spiral sulcus (ISS) along with the OoC. Periodic structures (orange) coupled the TM and OoC and gaps created fluid channels representing subtectorial spaces (STS) that joined the ISS and SV. Scale bar 50 *μ*m. B) The place-frequency map was nearly the same as the previous OoC-only model. C-F) Measurement locations for velocity, displacement, and pressure were identical to those in the pseudo-OoC model. E,F) Experimental ST pressure re. ear canal pressure (x-marks, Dong and Olson, 2013) and BM displacement re. ear canal pressure (Meenderink et al., 2022) are shown normalized to BF (Dong and Olson 90 dB in vivo BF 16 kHz, Meenderink 50 dB in vivo BF 2.2 kHz). G) Experimental wavenumber-frequency relationships re. BF are included from (De La Rochefoucauld and Olson, 2007)^7^ and derived from the Meenderink et al. phase shown in (F) (DLR & O 80 dB in vivo BF 10 kHz). H) The complex impedance plot showed the mass effects of the pectinate zone, OoC, TM, ISC, and enclosed fluid (STS and ISS) each added to the last. I) The eigenmode analysis produced three eigenfrequencies in the range of interest, connected by order (1^st^ being the lowest frequency). Above around 500 Hz, the second eigenfrequency corresponds to the classic traveling wave.

The place map (Fig. 7B) was nearly unchanged compared to the pseudo-OoC model with the BF remaining 2.5 kHz. The partition stiffness was only 60% higher and the spatial velocity magnitude (Fig. 7C) and phase (Fig. 7D) were similar to the OoC case including the second, apical wave. The displacement and pressure gain (Fig. 7E,F) told a similar story to the previous studies remaining stiffness-dominated. The model BM displacement gain, increasing near 10 dB/octave and peaking around 200 nm/Pa was very similar to that measured experimentally in the gerbil middle turn (Meenderink et al., 2022). Given the measurement was at 50 dB in vivo though, one might expect a more sharply tuned experimental response or conversely, a more broadly tuned postmortem model response. The phase slope was also similar, but values were offset around 0.5 cycles, which could be due to the measurement recording direction. The experimentally measured ST pressure gain (Dong and Olson, 2013) was also similar to the model pressure difference (and model ST pressure, not shown). The higher experimental magnitude could be due to the more basal measurement location (16 kHz BF instead of 2.5 kHz). Despite these differences, the model responses were largely comparable to the experimental results.

The dispersion of the full-length model was shown as dots in Fig. 7G-I as in the previous figures. The BF corresponded to a wavenumber around ¾ cycles/mm, similar to all previous studies and to experimentally obtained wavenumbers relative to BF (open circles (Meenderink et al., 2022) and black line (De La Rochefoucauld and Olson, 2007)). The wavenumber from de la Rochefoucauld and Olson was reported directly while the Meenderink et al. wavenumber was derived from the phase shown in (Fig. 7F)^8^. Each experimental wavenumber reached near 1.1 cyc/mm at BF, slightly greater than the 0.75 cycles/mm of the model. However, even this slight difference could be an overestimate as the data from de la Rochefoucauld and Olson was obtained at a higher frequency place and the Meenderink et al. data was from a similar frequency place but at low level in vivo. Figure 7H showed that using the 2D WKB approximation with a complex partition impedance, the model dispersion relation was most closely captured when considering the mass of the tissues and enclosed fluids.

The eigenfrequency analysis (Fig. 7I) now showed three modes: the slow traveling wave mode that overlapped well with the full-length model (predominantly 2^nd^ eigenfrequency (ef)), the faster compressive mode also seen in the OoC modification, and an additional mode tied to motion of the TM (predominantly 1^st^ ef). The three modes are distinct in their displacement and pressure profiles (Fig. 8). The classical traveling ‘slow wave’ was shown to have a pressure difference across the partition (Fig. 8E) which resulted in the entire partition moving in unison deflecting towards one scala (Fig. 8D). Since the analysis is harmonic, the motion would deflect towards the other scala and back again within one cycle. In the compressive ‘fast wave’ mode, the pressure was uniform across both scalae (Fig. 8G) and the BM and TM moved in opposite phases (Fig. 8F). Lastly, there was the ‘TM/ISS’ mode, labeled as such because the TM was the primary structure with a nonzero displacement (Fig. 8B) arising from pressure in the ISS (Fig. 8C). Here, the TM acted like its own partition secured at two ends with displacement maximal in the center. The pressure distribution corresponding to this motion showed the pressure difference across the SV and ISS and was entirely independent of ST. The high pressure in the ISS and low pressure above the TM lead to an upwards TM deflection. At low frequencies, the STS would vent the ISS rendering this pressure distribution impossible. Hence, such a mode had a low frequency limit (Fig. 8A inset, ∼500 Hz). This contrasts with the traveling wave which theoretically approaches zero in wavenumber (i.e. approaches infinite wavelength) as frequency goes to zero. Taken together, these require that a region of overlap existed between the two, as seen near 500 Hz.

**FIG 8.**
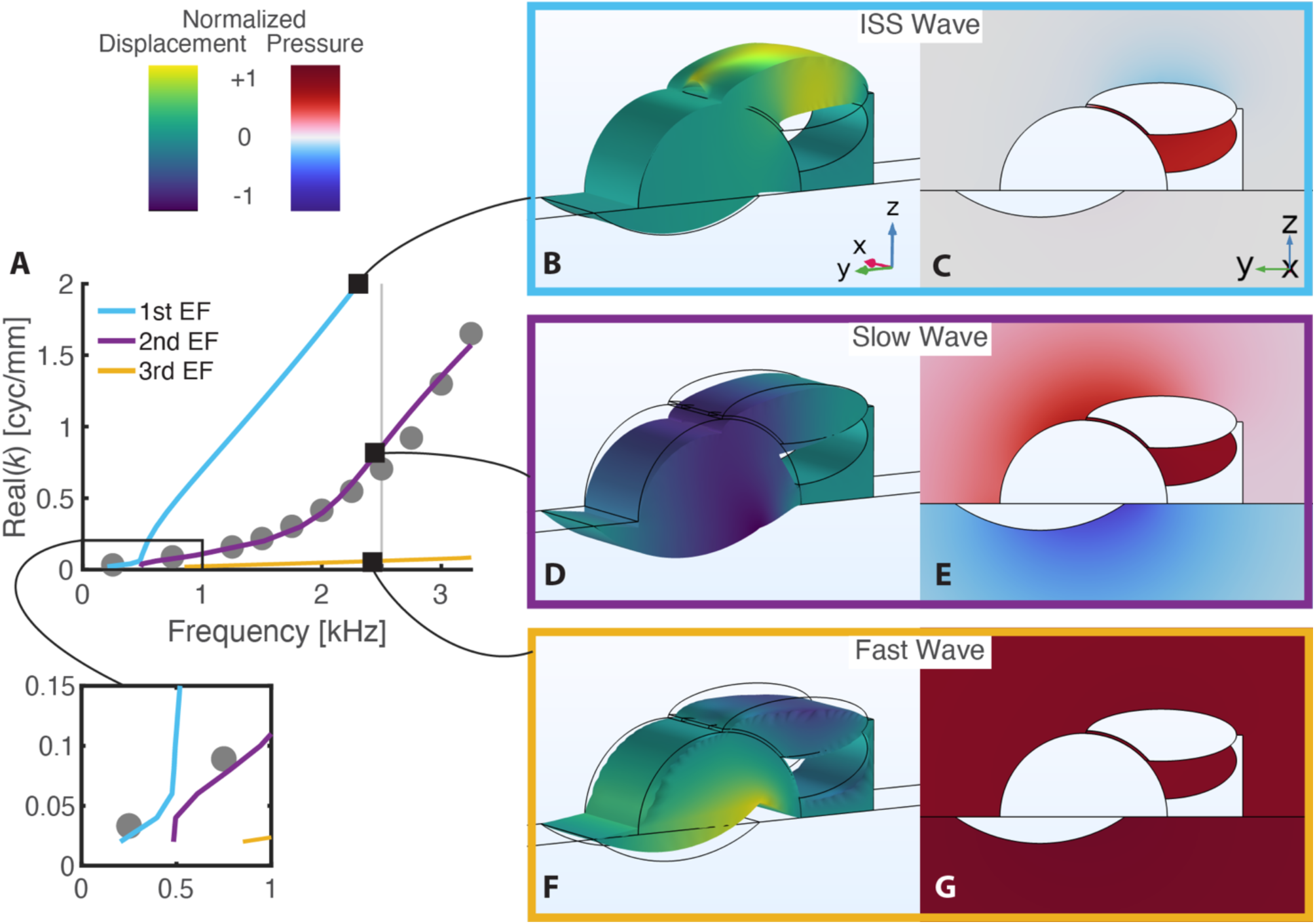
(Color online). Three modes of the TM, ISS model near the BF. A) The dispersion diagram from Fig. 7I with zoomed-in view of the modes at low frequencies (0-1 kHz). Lines are grouped by frequency (e.g. lowest frequency mode is light blue) and not by mode shape. The slow wave mode begins as the lowest frequency mode (light blue) before becoming the second lowest (purple). Black square markers indicate the wavenumber-frequency combination where each mode shape in B-G was captured. B-G) Displacement (B, D, F) and pressure (C, E, G) mode shapes. Color indicates normalized real transverse displacement (negative in dark blue to positive in yellow with zero in green) or pressure (negative in dark blue to positive in red with zero in gray). Eigenvalue problems do not produce absolute magnitudes of motion, so displacement scale factors were chosen to illustrate relative motions between structures. B,C) Displacement and pressure modes of the ‘ISS’ wave (k=2 cyc/mm, f=2306 Hz). The TM transverse motion occurs from a pressure difference between the ISS and SV without regard for ST pressure. D,E) Modes of the classic ‘slow’ traveling wave mode (k=0.8 cyc/mm, f=2445 Hz) with its simple bending displacement and pressure difference between the two scalae. Near the best frequency, pressure is focused at the partition and decays away as seen by the fading color towards the edges of the image. F,G) The ‘fast’ wave compressive mode (k=0.06 cyc/mm, f=2458 Hz) with in-phase high pressure in each scala squeezing the partition together.

## V. DISCUSSION

Full-length cochlear models have been used to explore a number of phenomena including frequency tuning (Steele and Taber, 1979; De Boer, 1981; Neely, 1985; Watts, 2000; Xia et al., 2018), the origins of amplification (Yoon et al., 2011; Wang et al., 2016), dynamic range (Sisto et al., 2021), and the impacts of cochlear implants (Elliott et al., 2011). However, while these models inherently produced traveling waves, they often considered the simplest morphological partition of a flat BM. Some finite-element models incorporated more intricate partitions with a tectorial membrane, reticular lamina, hair cells, and Deiters’ cells (Kolston and Ashmore, 1996; Motallebzadeh et al., 2018), but they did not explore how traveling wave dispersion, a property fundamental to the cochlea’s frequency selectivity, related to those morphological features.

Cochlear slice models offer an opportunity to investigate micromechanical interactions in the organ of Corti with the tradeoff that they do not produce the macromechanical signatures of full-length models. Several finite-element slice models have been developed with varying dimensionality, degrees of longitudinal coupling, and approaches regarding the traveling wave. Some did not incorporate a wavenumber-frequency relationship (Ni et al., 2016; Maftoon et al., 2018; Zagadou et al., 2020), others adjusted material parameters to fit data obtained experimentally (Cai and Chadwick, 2003; Cai et al., 2004), and some considered multiple modes of wave propagation beyond the classical traveling wave (Steele, 1999; Elliott et al., 2013).

Although computational models are well-suited for evaluating incremental changes to morphological features and have been used as a ‘numerical genetics laboratory’ (Motallebzadeh et al., 2018), such an evaluation had not been conducted for the traveling wave. In fact, an experimental investigation by Eze and Olson (2011) provided the closest comparison. As expected, controlling and evaluating the degree of tissue damage was a notable challenge. This investigation sought to fill that gap and explore how traveling wave dispersion and partition motion in the passive cochlea was affected by incremental variations of cochlear partition morphology. To bridge investigations of macro- and micromechanics, we also explored whether and how those effects could be predicted using a finite-element cochlear slice model.

### A. Key Findings & Comparison to Literature

A key finding was that upon additions of partition morphology in the full-length models, the BF increased roughly 2.5 octaves (Fig. 9F). Additionally, relative to BF, the displacement gain decreased and local pressure gain increased while there was no frequency dependence between the two (Fig. 9D). These suggested a prominent change in effective partition stiffness, which we quantified when evaluating the static deformation of the slice models. As for the traveling wave dispersion, we found it minimally varied relative to the BF (9H). To simulate wave propagation in the slice models, we incorporated Floquet boundary conditions and found the classical traveling wave dispersion was recovered using eigenfrequency studies (I in Figs. 4–7). Furthermore, the method produced a new dispersion curve within the frequency range of interest for the TM-ISS model which was associated with pressure in the ISS and motion of the TM (Fig. 7B,C). This mode of motion appeared to correspond with motion recently observed by Page et al. (2025, Supplement Movie 2). Importantly, we found this mode appeared without disrupting macromechanical features like the cochlear map, tuning, or the dispersion of the classical traveling wave (OoC versus TM in Fig. 9G,H).

**FIG. 9.**
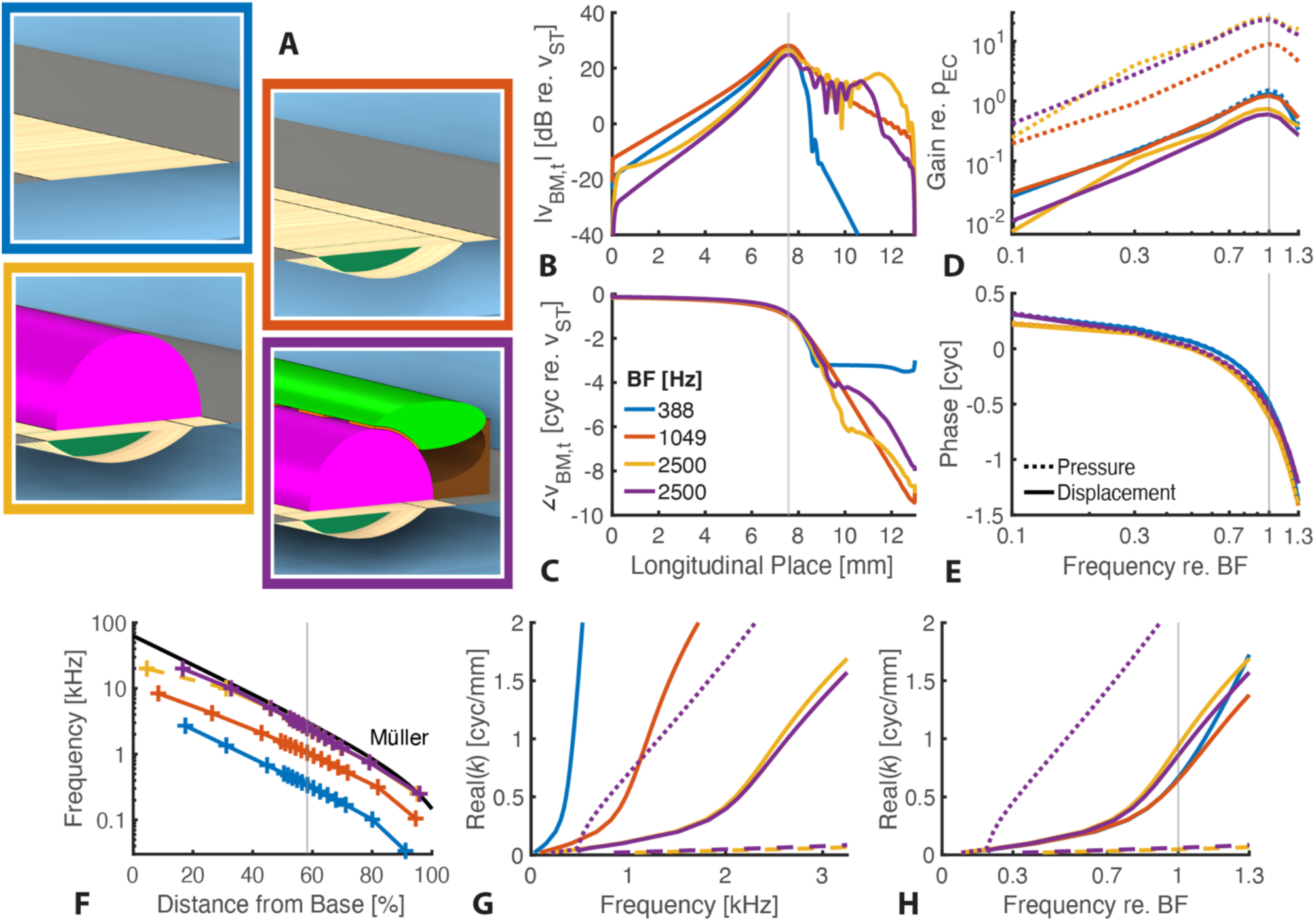
(Color Online). Comparing results across morphological variations. A) The four morphological variations studied. Line colors in each subsequent panel correspond to the border outlining each morphological variation (i.e. not the color of a particular structure): Flat BM (blue), Arch BM (orange), OoC (yellow), TM/ISS (purple). B,C) Normalized spatial velocity response magnitude (B) and phase (C) for the BF in each respective full-length model. Stimulation frequencies were chosen to peak at the same location. D,E) Displacement (solid lines) and local pressure (dotted lines) gain magnitudes (D) and phases (E) for each variation plotted relative to each model’s BF. Displacements were measured on the BM and local pressure differences were measured 1 *μ*m from the partition towards the ST and SV with partition locations as follows: BM center (Flat BM), arcuate-pectinate junction (APJ) (Arch BM), APJ for ST and OoC for SV (OoC & TM). Displacement phase was shifted 0.5 cycles so a positive pressure difference produced a positive displacement. F) Cochlear map across models with reference to the in vivo Müller map (black) (Müller, 1996). The vertical line indicated the longitudinal location for analysis which was unchanged across variations. G,H) Dispersion relation for each variation plotted against absolute frequency (G) and frequency relative to each BF (H) using the dispersion results from the eigenfrequency analyses. Solid lines predominantly represent the classical traveling wave. Above 0.2 times the BF, the dashed and dotted lines represent the compressive and TM/ISS modes, respectively.

#### 1. Stiffness Effects Dominate Local Displacement and Pressure

The displacement gain for the TM/ISS variation—peaking around 200 nm/Pa—was close to that measured experimentally in the gerbil middle turn (Meenderink et al., 2022). The local cochlear partition pressure difference was also measured experimentally to produce a maximum gain at BF around 26 dB (Kale and Olson, 2015), similar to our observations for the TM variation. For all changes in morphology, no frequency dependence was detected between the pressure and displacement indicating a stiffness-dominated response, which has also been shown experimentally (Dong and Olson, 2013). However, due to the experimental challenges of recording cochlear pressures and controlling tissue damage, no study to our knowledge has examined how local pressure changes with progressive morphological changes. We found the pressure could be accounted for entirely by the change in cochlear input impedance and middle ear admittance with frequency. This was because the local pressure difference relative to oval window pressure was unchanged with frequency as the wave propagated to the best place before decaying.

In a study examining neomycin-damaged cochleae, Eze and Olson (2011) reported a maximal downward shift in characteristic frequency of 0.5 octaves measured by phase accumulation. In contrast, we found the BF changed much more, close to 2.5 octaves across the most extreme morphological conditions tested, which approximates the largest morphological change of Eze and Olson. They attributed the frequency shift to a change in partition stiffness. Since the BM fibers remained in their prep, this meant the structures surrounding the BM fibers like the OoC only contributed to a 0.5 octave increase in BF. Our much larger BF shift suggests that the stiffness of the tissues surrounding the BM fibers play a much greater role in generating the effective partition stiffness than was previously described. We suspect their results were due to a combination of a difference in measurement location (∼20 kHz vs 2.5 kHz) and experimental challenges including systematically damaging tissue, measuring smooth tuning curves, and measuring from consistent longitudinal places.

Using the final iteration (TM, ISS model) as the closest approximation to a healthy partition measured by experimentalists, we found the stiffness was 3.2 × 10^11^ Pa/m^2^ (7.4× 10^6^ dyne/cm^3^ with 231 μm BM width) for a BF of 2.5 kHz. Taking the stiffness fit from Emadi et al. (2004) for our place of interest 58% from the base (0.124 N/m) and a beam theory conversion previously described (De La Rochefoucauld and Olson, 2007), this translated to 3.5 × 10^11^ Pa/m^2^ (8 × 10^6^ dyne/cm^3^) in the clamped condition—remarkably similar to our results. How this stiffness changes with changing partition morphology has not been as easily evaluated due to experimental challenges in tissue examination and removal. Because we were able to measure the model stiffness directly, we found that along with a factor of 6.4 increase in BF (∼2.5 octaves) there was a factor of 41 increase in stiffness. A resonance-based place map predicts a change in BF equal to the square root of the change in stiffness. Although the stiffness increase between morphological variations did not completely align with the BF increase expected for a resonance-based place map, the overall change in BF was in line with this mapping.

#### 2. Traveling Wave Dispersion in Full-length Models

A primary impetus to this work was to explore the traveling wave dispersion relationship and how it varied when incorporating morphological details of the partition. We found little change in the wavenumber relative to BF across variations. Quantitatively, the wavenumber increased linearly at low frequencies before increasing more sharply at higher frequencies reaching around ¾ cycles/mm at the BF or a wavelength of 1.3 mm. Similar values were extracted from experimental measurements in gerbil (Zosuls et al., 2021; Meenderink et al., 2022) with more basal places breaking scaling symmetry and having slightly sharper increases and higher wavenumbers through the BF (Van Der Heijden and Cooper, 2018).

The influence of partition parameters and fluid coupling on the wavenumber were less established. The dominant features governing wave propagation in the passive cochlea could be assumed to be stiffness, as indicated by the shift in BF across morphological variations, and scalae geometry, given the consistency of the wavenumber relative to BF since the scalae did not change between studies. While we found an effective mass from 3D fluid coupling was required to capture the dispersion curve in the flat BM study (Fig. 4G), the same was not true for the more detailed morphological studies (H in Figs. 5–7). There, the dispersion curves were well matched using a 2D coupling approximation and the mass of the tissues and enclosed fluid, like in the inner spiral sulcus.

Cochlear partition tissue and inner fluid mass was said to be required for capturing experimentally measured wavenumbers in the gerbil base but using a 3D fluid coupling calculation (De La Rochefoucauld and Olson, 2007)—a dimensional effect that has been found to play a larger role in the base (Altoè, 2026). Additionally, De La Rochefoucauld and Olson noted the impact of local resonance decreased apically, which meant that even though the physical mass of the partition increases towards the apex, this was not required to fit the data. It was unclear, though, whether the authors updated the fraction of scala width occupied by the BM in their analysis of more apical locations. Our findings point to a reinterpretation of their conclusions: the mass of the tissues and enclosed fluids remains the effective mass required to match experimental wavenumber-frequency relationships throughout the cochlea, but only a 2D coupling formulation is needed in more apical regions. One caveat is that if there are no tissues or enclosed fluid, as in the case of the flat BM model, 3D longitudinal fluid coupling is required.

In summary, we found that for a given place along the cochlear length, adding morphological features to the cochlear partition changed the stiffness and BF, but did not change the traveling wave dispersion curve relative to the BF. Additionally, this dispersion curve was best predicted across morphological variations when considering 2D longitudinal fluid coupling and the mass of the tissues and enclosed fluids.

#### 3. Traveling Wave Dispersion Recovered in Floquet Slice Models

Floquet theory has infrequently been invoked in cochlear mechanics literature (Kim and Yoon, 2024; Ko and Stockie, 2015) and not for the use case presented here aside from exploratory work (Zagadou, 2010; Wang and Puria, 2025). Eigenfrequency analysis of the cochlea, on the other hand, has been evaluated on several occasions. Steele (1999) used the WKB method to calculate propagation modes in a shell of revolution model of the Pakistani water buffalo. While results were not directly compared to experimental wavenumber findings or a full-length model, he found, like us, one mode that corresponded to the normal mode of propagation with about equal pressure and fluid displacement in each scala and which showed a typical steepening of the wavenumber with respect to place after linear growth more basal.

Cai and colleagues (Cai and Chadwick, 2003; Cai et al., 2004) similarly used the WKB method but restricted their modal search to only the classical forward-running wave where they instead used the eigenfrequency analysis to calibrate the material properties of their partition components such that the resulting dispersion relationship was similar to experimental results. The wave finite element approach in Elliot et al. (2013) (similar to flat BM variation here) showed that the primary wave or “wave 1” in their terminology was the dominant mode when compared to a full-length finite element model until the wave’s magnitude was small apical to the BF. Our results (Fig. 3H) supported the notion that wave 1 is the primary influence on the traveling wave, however we did not detect a higher order mode that dominated the response at higher frequencies.

To conclude, we established a reliable method of computing the traveling wave dispersion curve for a 3D cochlear slice model by performing an eigenfrequency study of the model with Floquet boundary conditions.

#### 4. Floquet Slice Models Produce Multiple Propagating Modes

Several studies examined propagation modes in the cochlea involving the TM and/or ISS, and although earlier work suggested there was one dominant forward going wave despite these added components (Chadwick et al., 1996; Chadwick, 1998), later work showed how multiple modes could propagate in the OoC (Steele, 1999; Steele and Lim, 1999; Lamb and Chadwick, 2011). While our results also showed that the OoC sustained multiple modes, the nature of the ISS activity differed. Steele and Lim (1999) reported high pressure in the sulcus occurred with high BM displacement in an active case representing the typical wave analyzed in single-mode models, which agreed with earlier findings by de Boer (1993). Additionally, a second mode with high pressure in the sulcus that propagated apical to the BF was detected, although the experimental impact of this mode was unclear. Steele (1999) similarly reported an ISS wave which propagated apical to the BF with relatively smaller wavenumbers than the primary TW and no transition between the two. We found a fundamentally different relation where the ISS mode exhibited a cutoff frequency and transition with the TW mode. This could be due to the separation of each fluid space in the earlier studies (no off-diagonal elements or “leaking” between spaces) while the channels were fluid coupled in our study. Our configuration had the effect of equalizing the pressure across the spaces at low frequencies where the STS acted as a vent to the ISS, which meant the ISS mode would only appear at frequencies high enough for the STS impedance to restrict flow between the ISS and scalae cavities.

Lamb and Chadwick (2011) on the other hand computed a TM mode which peaked basal to the basilar membrane best place. However, they evaluated a lumped-parameter model which placed the TM and BM in series without consideration of a sulcus. Interestingly, the basal tuning of the TM wave meant there was a point where mode conversion could occur as the wavenumbers of each mode overlapped. It has been argued this phenomena was responsible for frequency selectivity (Van Der Heijden, 2014) and that it may be present in experimental measurements at low levels in vivo (Van Der Heijden and Cooper, 2018). The phenomena was also present in a passive model with a pseudo OoC and an asymmetric tunnel of Corti (Wang and Puria, 2025). While we observed a transition frequency where wavenumbers between modes overlapped (∼500 Hz in Fig. 6H), our full-length model showed no deviation from the primary traveling wave mode with opposing pressures in the scalae and in phase motion across all structures (“parallel” to use Van der Heijden’s terminology). This meant the ISS mode interacted with the TW mode at the transition frequency but never dominated in a way that the traveling wave could be described as having swapped mode shapes.

In summary, slice models can reveal different propagating modes of motion present at the location of the model. This includes a mode associated with high pressure in the ISS which deflects the TM transversely. Additional modes of motion are expected with the inclusion of additional fluid chambers.

### B. Limitations

As stated in the introduction, creating and solving models that span the length of the cochlea becomes prohibitively challenging with increasing morphological detail. Therefore, while we modeled the partition with more tissue components than other box models, the most detailed case still paled in comparison to nature’s construction. Although this prevented us from analyzing the specific effects of incorporating multiple hair cells or a tunnel or Corti, for instance, we were still able to provide a road map and build intuition for such manipulations with the simplifications of a pseudo-organ of Corti and fluid tunnel (ISS).

A similar limitation exists regarding slice models. Calculating the traveling wave dispersion of an arbitrary cochlear slice model using the eigenfrequency method will involve models not explicitly evaluated here. While these relationships will vary from the ones shown here and from one another, we expect these to be variations on a theme because of the established long- and short-wave nature of the traveling wave. Our work showed that the technique held for changes in stiffness, effective fluid mass, solid mass, and the addition of multiple propagating modes, which will likely arise for models with additional fluid tunnels and cavities. The shape and influence of these other modes on OoC micromechanics remains to be evaluated.

The TM/ISS mode in particular was shown to interact with the traveling wave below the BF due to the STS coupling the ISS and SV at low frequencies. While we found the traveling wave dispersion using inviscid fluid in the eigenfrequency method closely tracked that of the full-length model with viscosity, that wave was carried by the BM and scalae fluid without need or care for an STS. However, since the STS has been hypothesized to be a primary source of partition damping (Allen, 1980; Prodanovic et al., 2015), viscosity could influence the ISS mode. Our preliminary results including viscosity suggest the ISS mode is still present in the frequency range of interest and largely unchanged at high wavenumbers and frequencies (not shown). At low wavenumbers though, the ISS and SV decouple due to viscous drag through the STS, which eliminated the modal transition near 500 Hz seen in Fig. 8A. Steele (1999) expected that viscosity would dictate the response amplitude thereby making negligible the effects of modes below a certain wavelength (above a certain wavenumber) where the response peaks. A modal analysis could help determine the effects of this wave and how they are influenced by TM stiffness, ISS area, and STS dimensions including hair bundle morphology.

Across morphological variations, the partition occupied no more than 2% of the combined scalae area at the place of interest. While we found the local partition pressure difference did not vary beyond several dB within a factor of 1.3 times the BF, this could be a consequence of the marginal contribution of the partition area to the overall scala area. Likewise, the minor variations between the primary TW dispersion relative to BF could be impacted by a larger proportional partition area as the effective scala height would shrink. Changes in the features would impact prescribed passive tuning and the determination of BF in an arbitrary slice model, respectively. This is most relevant in the apex where the partition occupies a large fraction of the scala area and viscosity has been shown to play a greater role in shaping the response (Sasmal and Grosh, 2019). How the local pressure difference and dispersion in such a region could be affected is a subject for future investigations.

### C. Future Work

This study established the eigenfrequency method using Floquet boundary conditions to determine dispersion relationships in cochlear slice models. Future work in more detailed slice models can investigate what other modes exist, whether they interact with the traditional traveling wave mode, and if so, how. For example, it is hypothesized that IHCs are driven by fluid flow (Dallos et al., 1972; Russell and Sellick, 1983; Steele and Puria, 2005) potentially due to a change in STS height (Nowotny and Gummer, 2006; Wang et al., 2021) in addition to shearing between the TM and reticular lamina (ter Kuile, 1900). The ISS mode is expected to influence fluid flow around IHC bundles, so future studies could investigate whether it contributes to phenomena present in auditory nerve fiber responses but not BM responses like peak splitting (Kiang, 1990; Ruggero et al., 2000) and preferential synchrony to strong frequency components in complex sounds like formants in speech (Delgutte and Kiang, 1984; Delgutte, 1997). Separately, a new theory posits that cochlear amplification arises from OoC area changes (Altoè et al., 2022; Guinan, 2022). Future studies could investigate whether the OoC complex exhibits its own mode or modes of motion and their frequency dependence. Recent experiments have shown the reticular lamina is flexible (Cho and Puria, 2022), so researchers could also investigate the existence of a bending mode within the structure.

Our studies, along with those by Wang and Puria (2025), provide a foundation for future research using slice models to test hypotheses about cochlear micromechanics, such as hair bundle adaptation, reticular lamina flexibility, and TM mutations. OHC hair bundle adaptation has been hypothesized to set the correct phase for the cochlear amplifier (Kraut and Puria, 2024), and a slice model could be used to explore relevant parameters such as the adaptation time constant. The implications of having three rows of OHCs and the consequences of losing different rows are still unresolved, particularly in light of recent experimental data showing motion differences across the rows (Cho and Puria, 2022). Future work should investigate the contribution of the individual rows to cochlear amplification and whether flexibility in the structure facilitates high gain. Several genetic mutations known to cause hearing loss are linked to structural abnormalities in the TM (Cheatham et al., 2014; Lemons et al., 2019; Bullen et al., 2020), so a detailed slice model of the mouse cochlea could be used to characterize the mechanical consequences of such malformations. Overall, detailed cochlear slice models could be used both to probe microscopic interactions that cannot currently be studied in vivo and to predict effects that can be tested experimentally.

### D. Conclusions

In conclusion, this work demonstrated that across changes in partition morphology, the dispersion relation was unchanged relative to a variation’s best frequency. We also established the Floquet eigenfrequency method to calculate the dispersion of cochlear slice models. By increasing model complexity in steps, we also showed that the partition can support multiple propagating modes which are detected by our eigenfrequency method. Lastly, we showed how a slice model with a TM and ISS produced a novel mode of motion that moves the TM transversely and squeezes fluid through the ISS. These findings identified key relationships between cochlear partition structures and the traveling wave which can support the development of more detailed slice models for deeper investigations of organ of Corti passive and active micromechanics.

## ACKNOWLEDGMENTS

The authors would like to thank Andrew Tubelli, Yanli Wang, and Julius Kraut for help with COMSOL Multiphysics. We would also like to acknowledge Charles Steele who sent us on the journey behind these studies years ago and Karl Grosh who provided a key insight into the Floquet method. This work was supported by the National Institutes of Health (NIH) National Institute on Deafness and Other Communication Disorders (R01DC07910, F31DC021079, and Grant T32 DC000038, awarded to G. Géléoc, supporting Gabriel Alberts). The contents are solely the responsibility of the authors and do not necessarily represent the official views of NIH.

## AUTHOR DECLARATIONS

## DATA AVAILABILITY

The data that support the findings of this study will be openly available on the Harvard Dataverse at https://dataverse.harvard.edu/dataverse/Otobiomech or from the corresponding author upon reasonable request.

## FOOTNOTES

1 Damping has been made constant (Altoè and Charaziak, 2023; Altoè and Shera, 2020; Elliott et al., 2011), proportional to other model parameters (Ni et al., 2017), and more involved as a nonlinear function to capture viscous and amplification effects (Sisto et al., 2021). It is widely discussed in active mechanics since the region of amplification can be interpreted a region of negative damping. This study concerns passive mechanics and will not consider damping effects.

2 This assumption breaks down in the cochlear apex. See Supplemental Materials II.A.2.

3 Exploratory work including viscosity produced thermal and vorticity modes that were not of interest for this investigation and impeded our search for the acoustic modes of interest which could be compared to the full-length model.

4 While the wavenumber calculation was presented through 1.3 times the BF, model responses were collected through 8 times the BF (3 octaves) for a more complete accounting of the place map.

5 The cochlear input impedance phase was near 0.15 cycles. There was roughly -0.1 cycles difference between the phase of the oval window pressure and that of the pressure difference about the partition. While the pressure difference about the partition is typically in phase with the partition displacement, model orientation meant that a positive pressure produced a negative deflection (0.5 cycle phase difference). Finally, partition velocity phase is shifted up 0.25 cycles from displacement for harmonic signals. Summing these phases gives a BM relative to stapes velocity phase of -0.2 cycles. Displacement and pressure phases for the various transfer functions can be seen in Supplemental Material section IV.C.

6 Below the 2 kHz middle ear cutoff frequency, the middle ear admittance magnitude increased at 6 dB/octave and the phase was one quarter cycle, consistent with a stiffness-dominated middle ear response. Since converting from velocity to displacement for a harmonic signal means dividing by frequency (i.e. multiplying by -6 dB/octave), this offset the slope of the transfer function giving the BM displacement re. ear canal pressure a similar profile to the BM velocity re. stapes velocity.

7 BM responses relative to stapes motion were originally presented in (Cooper, 2000).

8 The wavenumber was derived from the frequency response phase according to 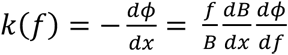 where *f* was the measured frequency, *B* was the BF of the measurement place, 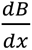 the slope of the frequency-place map about the measurement place, and 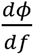 the frequency response phase.

