## Supplemental Information for "Cochlear wave dispersion reveals partition micromechanics"

Gabriel Alberts,[1,2] and Sunil Puria,[1,2,3]
*[1] PhD Program in Speech and Hearing Bioscience and Technology, Harvard University, Cambridge, MA 02138, USA*
*[2] Eaton-Peabody Laboratories, Massachusetts Eye and Ear, Boston, MA 02114, USA*
*[3] Department of Otolaryngology—Head and Neck Surgery, Harvard Medical School, Boston, MA 02115, USA*

### I. INTRODUCTION

The information presented here is to support the conclusions of the primary manuscript and expand on the results. The 'Methods' and 'Results' sections generally follow the structure of those sections in the primary text and include detailed information regarding geometry and material parameter choices in addition to physics governing equations in the finite-element models. While the primary text results were organized by morphological variation, we also explored certain topics across all variations. These are detailed in the 'Additional Results' section and include: radial profile of the basilar membrane, cochlear input impedance, calculating gain relative to ear canal pressure, calibrating a passive cochlear slice model, and three-dimensional longitudinal fluid coupling.

### II. METHODS

A full-length, finite-element box model of the gerbil cochlea was developed which contained numerous morphological features of the cochlear partition: an arch-beam representation of the basilar membrane (BM) (Kapuria et al., 2017; Xia et al., 2018), a solid mass above to represent a pseudo-organ of Corti (OoC), and a tectorial membrane (TM) complete with an inner spiral sulcus (ISS). The information provided here details the specific fits, modifications, and considerations that were made for each parameter as well as mesh and solver configurations. Additional subsections are included to discuss the construction of the Floquet slice model and modifications to the slice model for static stimulation of the partition.

#### A. Full-length Box Model

##### *1. Geometry*

A summary of length-dependent parameters with experimental values and model fits are shown in Fig. S1. BM width was linearly fit to data from the middle 80% of the cochlea from Plassmann et al. (1987), which was similar to values obtained by Schweitzer et al. (1996) (Fig. S1A). While not modeled explicitly in the 3D geometry, the BM fiber layers were implemented through the thickness of the 2D shell (see 'Physics' in the appendix of the main text). A logarithmic fit to the data in (Schweitzer et al., 1996) provided the decreasing thickness from base to apex of the upper and lower fiber layers of the BM (Fig. S1B), which were split around the pectinate zone (PZ) arch and combined in the arcuate zone. For the study not involving an arched BM (i.e. the traditional flat BM case), the summed upper and lower thicknesses was used for the entire BM width.

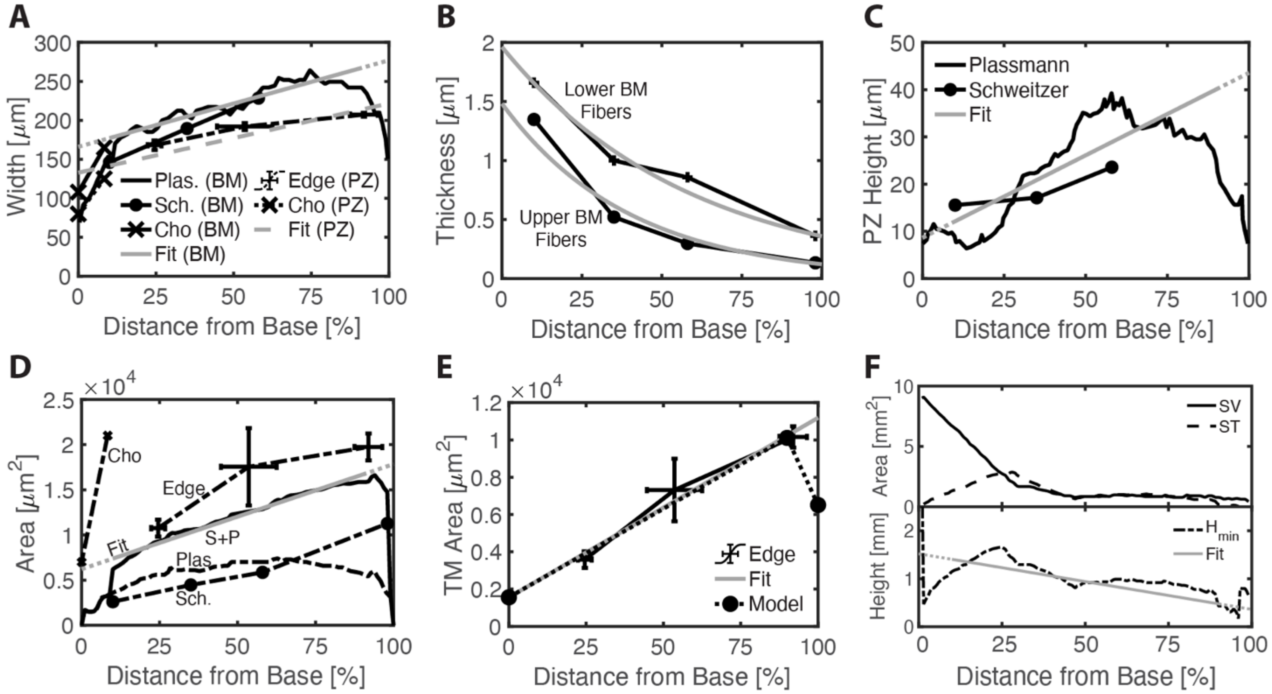


FIG. S1 Full-length model geometric parameters. All values plotted against percent distance from the base. Model values are equivalent to 'Fit' values except for TM area. Plassmann data was fit to the middle 80% of the cochlea (10-90% distance from base). A) BM, pectinate zone (PZ) width. BM width was made a linear fit to Plassmann and PZ width was set to 80% of BM width. B) BM fiber thickness. Fibers are split around the pectinate zone (PZ) arch and combined in the arcuate zone. 'Lower' refers to the fibers on the ST-facing side of the BM. Exponential fit to each layer. Data from Schweitzer. C) PZ arch height. Linear fit to Plassmann. D) Cho and Edge consider OoC area, Plassmann (Plas) reported Hensen cell and Claudius cell area, and Schweitzer (Sch) reported OoC area minus those lateral supporting cells (Hensen, Claudius). OoC area was determined from a linear fit to the sum of Plassmann and Schweitzer (S+P) from 10-90% distance. E) TM area. The fit was linear to the mean values of Edge. However, the TM was a complex shape whose area changed nonlinearly with distance. Values from the fit were chosen for the base and helicotrema (90% from base). Area at the apex was chosen for the TM to fit the diminished scala width without disrupting the rest of the model geometry. F) Model height was linearly fit to the square root of the smaller scala area between SV and ST (Plassmann). References: (Plassmann et al., 1987; Schweitzer et al., 1996; Edge et al., 1998; Cho et al., 2022).

The proportion of the arched PZ width to the width of the BM was reported to decrease slightly from approximately 95% in the base to 72% in the apex (Edge et al., 1998) although more recent measurements in the hook region found the ratio was near 75% (Cho et al., 2022). The model PZ width was simplified to a constant 80% of the BM width across the cochlear length. The PZ arch intersected with one edge of the BM and was modeled as a segment of a circle whose radius was determined by the PZ width and arch height. The height was determined from a linear fit to data from the middle 80% of the cochlea by Plassmann (1987), which was again similar to Schweitzer (1996) (Fig. S1C).

OoC area was determined from a linear fit to data from the middle 80% of the cochlea (Fig. S1D). The data in this case was the combined reported measurements of Hensen cell and Claudius cell area in Plassmann (1987) and OoC area minus those supporting cells in Schweitzer (1996), which were similar to other reported values of total OoC area including tissue and fluids (Edge et al., 1998). The OoC was constructed as a half cylinder with a radius dependent on distance from the base. The edge between the BM arcuate zone and bone served as one long edge of the OoC. Given the OoC area and cross-section shape of a semicircle, this meant the diameter was roughly three quarters the width of the BM, which fell in line with images and schematics from the associated references. While data from the hook region in vivo was similar to previously reported data at the very base, the area increased much more rapidly over the first millimeter (Cho et al., 2022). This was not considered here. Unpublished in vivo images using the same technology (optical coherence tomography) showed that near our place of interest (~60% from the base) in the gerbil middle turn the area could be roughly two to three times larger than the fitted value.

The TM was made by crafting an ellipse at the base (x=0), helicotrema (x=L-H), and apex (x=L) with its most radial endpoint (vertex) aligned to the top of the OoC at those three longitudinal distances. The 3D structure was formed automatically by connecting the surfaces from base to apex (lofting) such that the ellipse at the base gradually morphed into the ellipse at the helicotrema which again smoothly morphed to the apex. This generated an overlapping volume with the OoC which was removed from the TM in the final geometry. The TM was subdivided further to create the STS and pseudo-hair bundles. Specifically, the three μm parallel to the OoC which overlapped with the TM volume as this approximated the cilia bundle height near the 2 kHz place (Fettiplace, 2017). The major and minor axes of the TM were adjusted at the base, helicotrema, and apex to produce a length-dependent area in the final geometry that approximated reported measurements (Edge et al., 1998). Area was reduced from the start of the helicotrema to the apex since the wide, centered BM left little width for the off-centered TM. Due to the various modifications that shaped the final TM geometry, the area at a given distance from the base which isn't one of the three defined distances was slightly smaller than a linear interpolation of the area would suggest.

A similar process to the TM was used to create the ISS, which was bounded by the OoC, TM, and another structure that represented the inner sulcus cells and limbus. Although experimental data for ISS area is sparse unlike for the OoC and TM, the model ISS area was similar to data from the base (Cho et al., 2022) and increased with distance from the base at a similar rate to the TM area.

Longitudinally, the gap between the OoC and TM alternated periodically between solid and fluid sections. The periodic solid connections represented the connections from outer hair cell bundles, which are firmly embedded in the TM. The periodic subtectorial space (STS) fluid channels vented the ISS to the larger scala fluid. Since our goal was to create a corresponding slice model with a thickness well below the wavelengths of interest to reduce potential contamination from aliasing, the sections were chosen to alternate every 50 μm. This allowed us to create a slice model which included both the solid and fluid spaces that was 100 μm thick longitudinally—less than one-sixth of a wavelength for wavenumbers below 1.5 cycles per millimeter as were expected through the best frequency.

Scala area was determined by a fit to the smaller scala in Plassmann (1987) from the middle 80% of the cochlea, which kept the area above zero through the apex. The linear tapering in scala height and width was similar to Ni et al. (2017) with a smaller apical area that more closely approximated anatomical measurements. The helicotrema in mammals is formed by an opening in the bone modiolar to the OoC approximately in the apical 10% of the cochlear length (Wright and Roland, 2018). The helicotrema in our model was positioned adjacent to the BM pectinate zone (PZ) (lateral in a coiled cochlea), which kept its area constant across morphological variations. Since the opening was made by removing the bone between the PZ and scala wall in the apical 10%, the area was similar to areas found in similarly sized animals (Mountain et al., 2003).

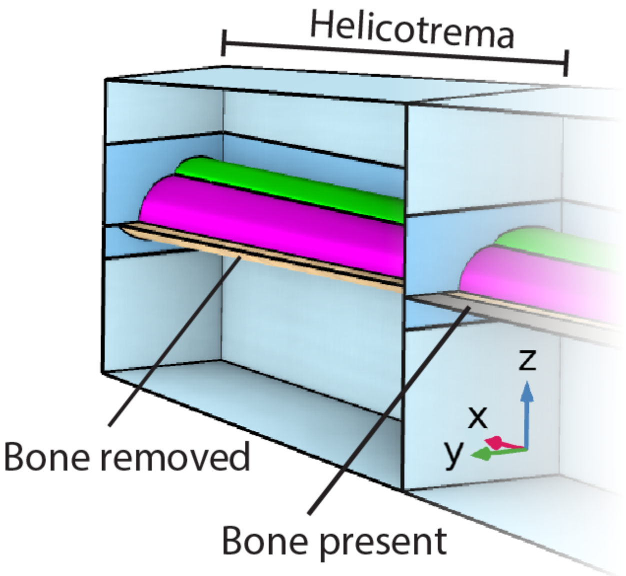

FIG. S2 Zoomed in view of helicotrema. The TM (green) and other structures occupy the "modiolar" side of the model (-y direction). For helicotrema consistency between morphological variations, bone was removed from the "lateral" side to connect the larger scala vestibuli and tympani. Here, gray bone is seen in the section basal (-x direction) to the helicotrema and is absent in the helicotrema.

#### *2. Materials*

The Poisson's ratio was 0.4 for the PZ, OoC, and TM to represent their incompressibility and was 0 for the BM to avoid unphysical stresses in shell elements (Motallebzadeh et al., 2018; "Structural Mechanics Module User's Guide," 2021). The bone attached to either side of the BM was modeled with fixed boundary conditions as the osseous spiral lamina and spiral ligament are typically considered immobile (Peterson and Bogert, 1950; Homer et al., 2004; Motallebzadeh et al., 2018). The 5% damping loss factor in the tissue was similar to previous investigations (Wang and Puria, 2025) and prevented unwanted sharp peaks and notches in the velocity response.

The author's best effort was made to apply an OoC Young's modulus which produced a frequency-place map near a half-octave below the in-vivo map from Müller (1996). This was most challenging in the apex where the best frequency was closer to or slightly higher than the in vivo best frequency. Viscosity is expected to lower the best frequency by reducing the response at higher frequencies and has been shown to play a greater role in tuning at the apex (Sasmal and Grosh, 2019). While included in our model, viscosity would be expected to have a larger effect on a more morphologically detailed model with numerous cells and fluid spaces in the OoC that greatly increase its surface area. However, several studies have also shown that passive tuning in the apex can be sharper with a higher best frequency than in the active case (Recio-Spinoso and Oghalai, 2017; Sasmal and Grosh, 2019; Recio-Spinoso et al., 2024). It is interesting then that despite a very compliant OoC, just the presence of our pseudo-OoC moved the best frequency up to that found in vivo. Since our studies were focused on the middle of the cochlea, these effects were not investigated further.

#### *3. Physics and Multiphysics Couplings*

The physics and coupling equations used in th1e models were derived from COMSOL Multiphysics modules ("Acoustics Module User's Guide," 2021; "Structural Mechanics Module User's Guide," 2021) and are presented here for completeness.

**Physics**

The thermoviscous acoustics domain solved the mass continuity equation and the linearized Navier-Stokes equation for acoustic perturbations of compressible Newtonian fluids, which for time-harmonic signals were of the form

$$i\omega\rho_f + \nabla \cdot (\rho_f \mathbf{u}_f) = 0 \quad (1a)$$

$$i\omega\rho_f \mathbf{u}_f = \nabla \cdot \left[ -p_{\mathrm{va}}\mathbf{I} + \mu\left(\nabla \mathbf{u}_f + (\nabla \mathbf{u}_f)^{\mathrm{T}}\right) - \left(\frac{2\mu}{3} - \mu_B\right)(\nabla \cdot \mathbf{u}_f)\mathbf{I}\right] \quad (1b)$$

with $i = \sqrt{-1}$, $\omega$ the radian frequency, $\rho_f$ the fluid density, $\nabla$ the del operator, and $\mathbf{u}_f$ the 3D fluid velocity. In addition, $p_{\mathrm{va}}$, $\mathbf{I}$, $\mu$, and $\mu_B$ were the viscoacoustic fluid pressure, identity matrix, dynamic viscosity, and bulk viscosity, respectively. The Fourier heat conduction law was also solved although temperature effects were not considered in our studies. Nonetheless, the domain consisted of three dependent variables: pressure $p_{\mathrm{va}}$, velocity $\mathbf{u}_f$, and temperature. Pressure was solved for with linear shape functions while velocity and temperature were solved using quadratic serendipity shape functions.

In the pressure acoustics formulation, the fluid was considered inviscid, which reduced the viscoacoustic equations above to an inhomogeneous Helmholtz equation

$$\nabla \cdot \left(-\frac{1}{\rho_f}(\nabla p_{\mathrm{pa}} - \mathbf{q}_{\mathrm{d}})\right) - \frac{\omega^2 p_{\mathrm{pa}}}{\rho_f c^2} = Q_m \quad . \quad (2)$$

Here, $\mathbf{q}_{\mathrm{d}}$ represented dipole sources of which there were none, and $Q_m$ was a source acceleration, only present as the time derivative of the harmonic prescribed velocity over the oval window area. Furthermore, $c$ was the speed of sound, and $p_{\mathrm{pa}}$ was the total acoustic pressure which equaled the scattered field pressure. Pressure was the only dependent variable in the domain and was implemented with quadratic Lagrange shape functions.

The equation of motion for solids in the frequency domain was derived from Newton's second law

$$-\rho_s \omega^2 \mathbf{x}_s = \nabla \cdot \mathrm{S_s} + \mathbf{F}e^{i\boldsymbol{\varphi}} \quad (3)$$

where $\rho_s$ was solid density, $\mathbf{x}_s$ 3D solid displacement, $\mathrm{S_s}$ the stress tensor, and $\mathbf{F}$ external force. All forces acting on the solid were applied via coupling with fluid and shell interfaces so $\mathbf{F} = \mathbf{0}$. All structures were linear elastic making the stress tensor the double contraction of the $4^{\mathrm{th}}$-order elasticity tensor and $2^{\mathrm{nd}}$ -order elastic strain tensor, modified by an isotropic loss factor $\eta$

$$\mathrm{S_s} = \mathbf{C}\colon \epsilon(1 + i\eta) \quad (4)$$

with

$$\epsilon = \frac{1}{2}((\nabla \mathbf{x}_s)^{\mathrm{T}} + \nabla \mathbf{x}_s) \quad (5)$$

and $\mathbf{C}$ a function of the Young's modulus and Poisson's ratio. Additionally, the strain tensor was a symmetric 3x3 matrix and the elasticity tensor was reduced to a symmetric 6x6 matrix. The analysis was geometrically linear, so the first and second Piola-Kirchhoff stresses and Cauchy stress were equivalent, and the strain was the engineering strain. The three components of displacement served as the dependent variables, and elements were discretized using quadratic serendipity shape functions.

The BM fiber layers were modeled as shells to reduce computational cost for structures that exhibit significant bending stiffness and are thin compared to their width and length along the cochlea. Shell quantities were defined relative to their midsurface such that the equation of motion took the form

$$-\rho_{sh}\omega^2(\mathbf{x}_{sh} + \mathrm{z}\mathbf{a}) = \nabla \cdot \mathrm{S_{sh}} + \mathbf{F} + 6(\mathbf{M} \times \mathbf{n})\frac{z}{d} \quad (6)$$

with $\rho_{sh}$ representing shell density, $\mathbf{x}_{sh}$ shell displacement, $\mathbf{a}$ displacement of the shell normal, and $z$ the local thickness direction which varied from -1 at the shell's bottom surface to +1 at the top with 0 defined as the midsurface. With negligible transverse stress ($\sigma_z = 0$) assumed, integration through the thickness direction was performed analytically. Additionally, $\mathbf{S}_{\mathbf{sh}}$ was the shell stress resultant tensor, $\mathbf{F}$ and $\mathbf{M}$ the external force and moment vectors, respectively, $\mathbf{n}$ the shell normal, and $d$ the shell thickness. Under the plane stress assumption that underlies shell theory, strains and stresses were reduced and partitioned to represent membrane, bending, and shear action which were transformed separately and combined in the weak form. The membrane forces, bending moments, and shear forces were computed from the corresponding stresses as defined by

$$N = d\mathrm{S}_{\mathrm{sh,m}} \tag{7a}$$

$$M = \frac{d^2}{6}\mathrm{S}_{\mathrm{sh,b}} \tag{7b}$$

$$Q = d\mathrm{S}_{\mathrm{sh,s}} \tag{7c}$$

where

$$\mathrm{S}_{\mathrm{sh,m}} = \mathbf{C}\!:\gamma(1 + i\eta) \tag{8a}$$

$$\mathrm{S}_{\mathrm{sh,b}} = \frac{d}{2}\mathbf{C}\!:\chi(1 + i\eta) \tag{8b}$$

$$\mathrm{S}_{\mathrm{sh,s}} = \frac{5}{6}\mathbf{C}\!:\zeta(1 + i\eta) \tag{8c}$$

were the membrane, bending, and shear stresses, respectively. The in-plane stress could then be computed as $\mathrm{S}_{\mathrm{sh,p}} = \mathrm{S}_{\mathrm{sh,m}} + z\mathrm{S}_{\mathrm{sh,b}}$. The plane stress constitutive matrices $\mathbf{C}$ were a function of the Young's modulus and Poisson's ratio, and the corresponding membrane, bending, and shear strains $\gamma$, $\chi$, and $\zeta$ were defined for a geometrically linear analysis as

$$\gamma_{\alpha\beta} = \frac{1}{2}\left[\frac{\partial \mathbf{x}_{sh}}{\partial \xi^\alpha}\frac{\partial \mathbf{r}}{\partial \xi^\beta} + \frac{\partial \mathbf{r}}{\partial \xi^\alpha}\frac{\partial \mathbf{x}_{sh}}{\partial \xi^\beta} + \frac{\partial \mathbf{x}_{sh}}{\partial \xi^\alpha}\frac{\partial \mathbf{x}_{sh}}{\partial \xi^\beta}\right] \tag{9a}$$

$$\chi_{\alpha\beta} = \frac{1}{2}\left[\frac{\partial \mathbf{r}}{\partial \xi^\alpha}\frac{\partial \mathbf{a}}{\partial \xi^\beta} + \frac{\partial \mathbf{a}}{\partial \xi^\alpha}\frac{\partial \mathbf{r}}{\partial \xi^\beta} + \frac{\partial \mathbf{x}_{sh}}{\partial \xi^\alpha}\frac{\partial \mathbf{n}}{\partial \xi^\beta} + \frac{\partial \mathbf{n}}{\partial \xi^\alpha}\frac{\partial \mathbf{x}_{sh}}{\partial \xi^\beta}\right] \tag{9b}$$

$$\zeta_{\alpha\beta} = \frac{1}{2}\left[\frac{\partial \mathbf{r}}{\partial \xi^\alpha}\cdot \mathbf{a} + \frac{\partial \mathbf{x}_{sh}}{\partial \xi^\alpha}\cdot \mathbf{n}\right] \tag{9c}$$

where $\mathbf{r}$ was the undeformed position of the shell, $\alpha$ and $\beta$ were indices ranging from 1 to 2, and $\xi^{\alpha,\beta}$ were the local coordinates along the shell's midsurface. To appropriately couple each boundary and surrounding solid structures, all shell midsurfaces were centered (i.e. no offset in the thickness direction). Dependent variables for the shell domain were the components of both midsurface displacement and shell normal displacement and were solved for using quadratic shape functions.

**Multiphysics Coupling**

Continuity in normal acceleration and normal stress was enforced between the pressure and viscoacoustic domains by

$$-\mathbf{n}\cdot\left(-\frac{1}{\rho_f}\nabla p_{\mathrm{pa}}\right) = -\mathbf{n}\cdot i\omega\mathbf{u}_f \tag{10a}$$

$$\left[-p_{\mathrm{va}}\mathbf{I} + \mu\left(\nabla\mathbf{u}_f + (\nabla\mathbf{u}_f)^{\mathbf{T}}\right) - \left(\frac{2\mu}{3} - \mu_B\right)(\nabla\cdot\mathbf{u}_f)\mathbf{I}\right]\mathbf{n} = -p_{\mathrm{pa}}\mathbf{n} \tag{10b}$$

using the frequency domain formulation. Fluid-structure interfaces for both solid and shell domains enforced velocity continuity

$$\mathbf{u}_f = i\omega\mathbf{x}_{sh}^{s} \tag{11}$$

while displacement continuity was enforced between the two structural domains

$$\mathbf{x}_s = \mathbf{x}_{sh}\ . \tag{12}$$

***4. Mesh***

Modeling the BM fiber layers as 2D shells as opposed to 3D solids reduced the number of degrees of freedom (DoFs) for the fiber layers by at least 60% and avoided creating elements with extremely high aspect ratios—some over 150—given the small thickness of the layers. Given the structural similarity of the partition along the length of the model, we attempted to generate hexahedral and prism elements as often as possible with structurally similar cross-sectional meshes. The partition as a whole then maintained the same longitudinal resolution as the fluid STS channel of 12 nodes per 100 μm longitudinally (six quadratic elements), which was greater than the mesh density needed to resolve the shortest wavelengths of interest (12 nodes per 500 μm corresponding to a wavenumber of 2 cycles/mm). Scala fluids (i.e. excluding the ISS and STS fluids) were mesh after the partition as unstructured tetrahedral elements.

*5. Solver Configurations*

We used a linear, stationary solver for finding the steady state solution with automatic detection of matrix symmetry and sparsity in line with COMSOL default settings ("COMSOL Multiphysics Reference Manual," 2021). Constraint handling was done automatically such that nodal constraints were handled explicitly and one of the 'Orthonormal' or 'Sparse' methods was used for the remaining constraints. A fully coupled approach was taken using a damped Newton method with initial and minimum damping factors of 1 and 0.0001, respectively. The approach used a tolerance termination technique that allowed a maximum of 25 iterations and followed the "solution or residual" criteria where the nonlinear iterations terminated when the relative tolerance exceeded the relative error computed as the minimum of the solution-based error and the error given by the residual factor (1000) times the residual-based error. The simulations ran a PARDISO solver with automatic out-of-core memory allocation.

**B. Floquet Slice Model**

Generating a slice between two cross-sections of the full-length model at two longitudinal places would maintain some length-dependent parameters, so one cross-section was obtained at a single longitudinal place and its geometry copied across the thickness of the slice. The thickness was the smallest possible length that captured all the morphological features of the full-length model. The fluid STS and solid pseudo-hair bundles were the smallest features of interest and each extended 50 μm longitudinally one after the other. Therefore, the slice was made 100 μm thick to resolve one of each solid and fluid section as the alternating pattern would be computationally treated as repeating like in the full-length model. To avoid a physics mismatch across longitudinal faces, the fluid STS was placed in the center of the slice making the space between TM and OoC solid at each longitudinal face. The solution should be identical with the solid and fluid spaces inverted and was chosen for ease of visualization. Across the slice thickness, material properties were taken to be those from the longitudinal place of the original cross-section.

The mesh was required to be compatible between pairs of faces under the Floquet boundary condition, so the partition (including all solids and the fluid STS and ISS) was first constructed to have a dense mesh of hexahedral and prism elements with structurally identical cross-sectional meshes. The surrounding scala fluid was meshed with tetrahedral elements.

Eigenfrequency solutions were computed for a single prescribed Floquet wavenumber searching for four eigenfrequencies around 1000 Hz, closest in absolute value. The procedure was repeated for wavenumbers ranging from 0.02 to 0.1 cycles/mm in steps of 0.02 and 0.1 to 3 cycles/mm in steps of 0.1 using an ARPACK eigenfrequency solver and an automatic determination whether to use a real symmetric eigenvalue solver. The maximum number of iterations was kept at the default of 300. Likewise, Krylov dimensions were automatically approximately as twice the number of specified desired eigenfrequencies. Eigenvectors were RMS scaled, and matrix symmetry, sparsity, and constraint handling was done automatically as in the full-length model. The simulations ran a MUMPS solver with automatic out-of-core memory allocation.

**C. Volume Stiffness Calculation from Slice Model, Method 1**

To calculate the partition stiffness, the slice model was adapted for a static pressure load. The load, representing a pressure difference across the partition, was independent of the eigenfrequency study performed on the Floquet slice model and was not considered as a prestressed state for any subsequent dynamic analysis. Taking the fluid to be incompressible, as in Puria and Steele (2008), any transverse BM displacement would result in an equivalent scala fluid displacement. Therefore, all fluid was removed from the slice model and only the BM displacement was measured from an pressure load applied directly to the BM (Fig. S3). Longitudinal faces were unconstrainted as the study did not consider wave propagation. Stiffness was calculated as applied pressure divided by displaced traverse BM area.

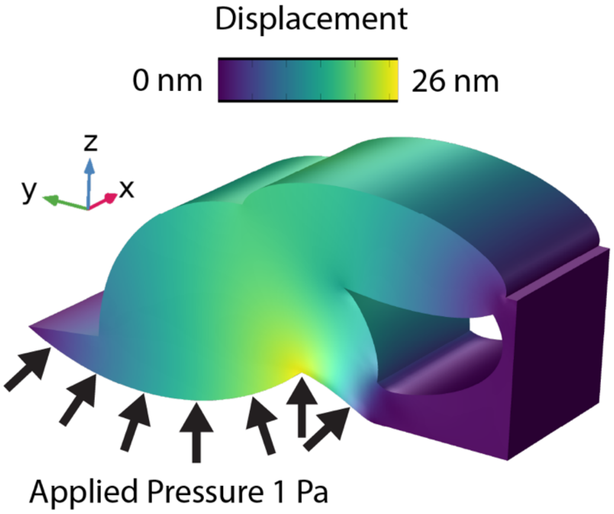


FIG. S3 Static displacement of TM model variation under a 1 Pa pressure load on the scala tympani-facing BM. Pressure is perpendicular to the surface (arrows) and produced a maximal displacement (yellow) at the junction between the arcuate and pectinate zones of the BM.

The solid and shell structures respectively now satisfied the equations

$$0 = \nabla \cdot \mathrm{S_s} \quad (13)$$

$$0 = \nabla \cdot \mathrm{S_{sh}} + \mathbf{F} + 6(\mathbf{M} \times \mathbf{n})\frac{z}{d} \quad (14)$$

where $\nabla$ is the del operator and $\mathrm{S}$ the stress tensor. Additionally, $\mathrm{S_{sh}}$ is the shell stress resultant tensor, $\mathbf{F}$ and $\mathbf{M}$ the external volumetric force and moment vectors, respectively, $\mathbf{n}$ the shell normal, $z$ the local thickness direction (-1 at the shell's bottom surface to +1 at the top with 0 defined as the midsurface), and $d$ the shell thickness. A 1 Pa pressure load was applied on the scala tympani-facing BM surface such that

$$\mathbf{F} = \mathrm{p}\mathbf{n}/\mathrm{d} \quad (15)$$

where $\mathrm{p}$ was the applied pressure and $\mathrm{d}$ the shell thickness resulting in units of force per volume. Structural components and materials were otherwise unchanged, and BM edges remained clamped. The mesh was generated by sweeping a quadrilateral mesh across the structure resulting in 117,579 DoFs. The same conditions were in place for detecting matrix symmetry and sparsity and handling constraints and out-of-core memory allocation. The simulation ran a MUMPS solver using a fully coupled approach that mirrored that from the full-length model. In all, the static deformation took 30 seconds to solve.

#### D. Finite Element Equations

##### Floquet Slice Model

Floquet boundaries related the dependent variables across a given physics according to a spatial periodicity independent of the geometric periodicity (Fig. 2, main text). For example, while the slice geometry was repeated every 100 μm (the slice thickness), the variables (e.g. pressure, velocity) could be spatially periodic over 1000 μm (1 cycle per millimeter). For solids, the displacement followed

$$\mathbf{x}_{\mathrm{dst}} = \mathbf{x}_{\mathrm{src}} e^{-i\mathbf{k}_F\cdot(\mathbf{r}_{\mathrm{dst}}-\mathbf{r}_{\mathrm{src}})} \quad (16)$$

Where $\mathbf{k}_F$ was the 3D Floquet wavenumber. For fluids, the pressure and pressure gradient were constrained by

$$p_{\mathrm{pa,dst}} = p_{\mathrm{pa,src}} e^{-i\mathbf{k}_F\cdot(\mathbf{r}_{\mathrm{dst}}-\mathbf{r}_{\mathrm{src}})}. \quad (17a)$$

$$-\mathbf{n}_{\mathrm{dst}}\cdot\left(-\frac{1}{\rho_f}\nabla p_{\mathrm{pa}}\right)_{\mathrm{dst}} = \mathbf{n}_{\mathrm{src}}\cdot\left(-\frac{1}{\rho_f}\nabla p_{\mathrm{pa}}\right)_{\mathrm{src}} e^{-i\mathbf{k}_F\cdot(\mathbf{r}_{\mathrm{dst}}-\mathbf{r}_{\mathrm{src}})}, \quad (17b)$$

Only the longitudinal direction had a non-zero wavenumber, so the dot products resulted in $\mathrm{k}_F\Delta x$ where $\Delta x$ was the slice thickness as shown in Fig. 2. The 2D shell fiber layers were not included in the Floquet boundary condition to avoid over constraining the model as the surrounding solid and fluid faces maintained their coupling to the shell layers.

The removal of viscoacoustic physics meant there was no longer coupling between two fluid physics, and fluid-structure coupling was to pressure acoustics as described mathematically by

$$-\mathbf{n}\cdot\left(-\frac{1}{\rho_f}\nabla p_{\mathrm{pa}}\right) = -\mathbf{n}\cdot\mathbf{u}_{\mathrm{tt}}, \quad (18a)$$

$$\mathbf{F}_{\mathrm{A}} = -p_{\mathrm{pa}}\mathbf{n} \quad (18b)$$

for exterior boundaries with $\mathbf{u}_{\mathrm{tt}}$ the structural acceleration and $\mathbf{F}_{\mathrm{A}}$ the load experienced by the structure in units of force per area. On interior boundaries that separate two adjacent fluid domains (i.e. shell between two fluids), the load was given by the pressure drop across the boundary and the structural acceleration experienced by the fluid was defined for each side of the boundary:

$$-\mathbf{n}\cdot\left(-\frac{1}{\rho_f}\nabla p_{\mathrm{pa}}\right)_{\mathrm{up,down}} = -\mathbf{n}\cdot\mathbf{u}_{\mathrm{tt}}, \quad (19a)$$

$$\mathbf{F}_{\mathrm{A}} = (p_{\mathrm{pa,down}} - p_{\mathrm{pa,up}})\mathbf{n}. \quad (19b)$$

### III. RESULTS

#### A. Flat BM

The model displayed a magnitude notch apical to the peak, most clearly visible in the orange line of Fig. 4C of the main text which peaks around the 10 mm place. The notch was similar to those first observed by Rhode (1971) and has been attributed to interaction between the fast fluid compression wave and slow traveling wave (Cooper and Rhode, 1996; Recio-Spinoso and Rhode, 2015) a second vibrational mode (Watts, 2000), and reflections due to finite scala height (Zweig, 2015). The phase responses in Fig. 4D showed the characteristic traveling wave phase accumulation, which decreased linearly at first then at an increasingly faster rate[1] after roughly one cycle consistent with passive or postmortem experiments in gerbil (Altoè and Shera, 2020; Cho and Puria, 2022; De La Rochefoucauld and Olson, 2007). Additionally, the magnitude notch aligned with a notch in phase and slight increase to a plateau. Plateaus between frequencies were separated by close to an integer number of cycles with small differences potentially due to interaction with reflections. While Zweig's proposal considered reflections, it hypothesized a slight plateau before a sharp decrease to a new phase plateau. There was also no vibrational mode which appeared to interact with the primary mode. Therefore, these data were most consistent with the influence of the fast compression wave.

#### B. Arch BM

Compared to the flat BM, the introduction of an arched pectinate zone produced a similar velocity slope approaching the peak but notably reduced the slope apical to the best place from -60 dB/mm to -12 dB/mm just past the peak (Fig. 5C). Despite the larger partition response apical to the best place, the peak velocity was nearly the same as the flat BM model. For lower frequencies, the magnitude and phase displayed ripples near the apex indicative of reflections from the helicotrema. The velocity peak was reached after about one cycle of phase, like the flat BM model, although the phase only plateaued for the two highest frequencies tested and reached ten cycles of phase accumulation for 1.3 times the BF. The larger wave amplitude and continued phase accumulation represented the dominance of the traveling wave mode apical to the peak.

The wavenumber of the full-length model increased linearly at first before turning quadratic in frequency in the short-wave region, as predicted by the 2D approximation based solely on the partition stiffness (Fig. 5G). The effect of the added mass of the pectinate zone ground substance was to further increase the wavenumber in the short-wave region (Fig. 5H, dark line, $m_{PZ}$). While this increase was slightly beyond what was measured in the full-length arch BM model, the slope of the dispersion curve including the ground substance mass was more similar to the full-length model than the massless case (m=0). Decreasing the partition stiffness by ~20% provided a near overlap through BF (not shown). It is possible that the effects of the added mass were modified by its asymmetric nature and enclosure within the BM fiber layers.

Partition displacement and velocity were measured at the junction between the arcuate and pectinate zones, although the responses were qualitatively similar when measured at other radial locations. The local pressure difference was measured 1 μm above and below the junction between the zones, but this was also similar when measured at other radial locations.

### IV. ADDITIONAL RESULTS

The results presented here are organized by topic and consider all four morphological variations examined in the main text: flat BM, arch BM, pseudo-OoC, and TM/ISS.

#### C. Radial Profile

The introduction of the pectinate zone arch broke the radial and transverse symmetry of the flat BM box model. The effect was prominent in the radial profile of the BM displacement, which went from approximately a square cosine function peaking in the center for the flat BM variation (a plate clamped on two ends) to a skewed profile peaking at the junction between the arcuate and pectinate zones (Fig. S4). This profile was maintained across frequencies up to twice the BF of a given variation[2] and through the addition of masses atop the BM in the OoC and TM variations.

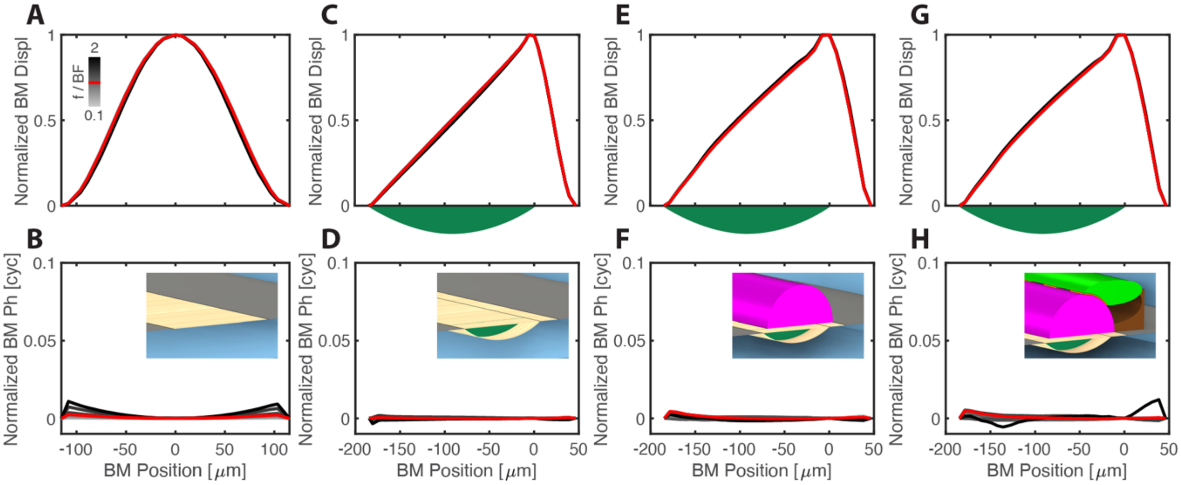


FIG. S4. (Color online) BM radial profile across full-length model variations. Profile magnitude (top row) and phase (bottom row) presented for four partition morphologies (columns). Magnitude normalized to peak displacement at each frequency. Phase normalized to phase at BM position zero (radial center for left column Flat BM variation and arcuate-pectinate junction for other variations/columns) with radial edges set to zero phase. This removed phase differences between frequencies due to the traveling wave. Frequencies ranged from 0.1 to 2 times the BF for each variation as indicated by the grayscale bar in (A) with the BF response in red. Insets in phase plots show morphology for the variation corresponding to that column. A,B) Flat BM variation magnitude (A) and phase (B) showed a symmetric radial profile with maximal displacement in the center and near uniform phase across the width of the BM. C,D) Arch BM variation magnitude (C) and phase (D) with data from the flat, SV-facing side of the BM—the same boundary as in the Flat BM variation. The magnitude profile showed a distinct asymmetry that peaked at the end of the pectinate zone arch. E,F) OoC variation magnitude (E) and phase (F) showed a similar asymmetric profile as the Arch BM variation with slightly more motion in the pectinate zone and still no phase difference across the BM width. G,H) TM variation magnitude (G) and phase (H) showed maintained radial asymmetry and consistent phase.

The arched BM model construction followed Kapuria et al. (2017) and Xia et al. (2018), which showed that the arch made for a stiffer pectinate zone that better matched the radial profile measured by Cooper (2000) and Cooper and Van der Heijden (2018). Previous investigations evaluated both hinged and clamped boundary conditions (Elliott et al., 2010; Homer et al., 2004; Steele and Taber, 1979). While Homer concluded a combination of the two boundary conditions was necessary to replicate the radial profile, we show the profile emerges from the morphology of the gerbil basilar membrane which imposes a natural boundary condition that is neither fixed nor hinged.

#### D. Cochlear Input Impedance

We examined the effects of incremental adaptations to the cochlear partition morphology on responses at one longitudinal place like the displacement, pressure difference, and dispersion. However, those adaptations were implemented across the full length of the cochlear model with length-dependent changes in geometry and material properties. The whole-cochlea effects of such modifications could be captured by the cochlear input impedance, commonly defined as

$$Z_C = {}^{p_{OW}}/_{U_{ST}} \tag{20}$$

with $p_{OW}$ the oval window pressure as measured in scala vestibuli and $U_{ST}$ the stapes volume velocity in the piston direction (equivalent to longitudinal direction, x) or the linear velocity multiplied by the 0.62 mm$^2$ footplate area. Numerous experiments have characterized the input impedance of gerbil, a common laboratory animal. Our results are compared to these data in Fig. S5.

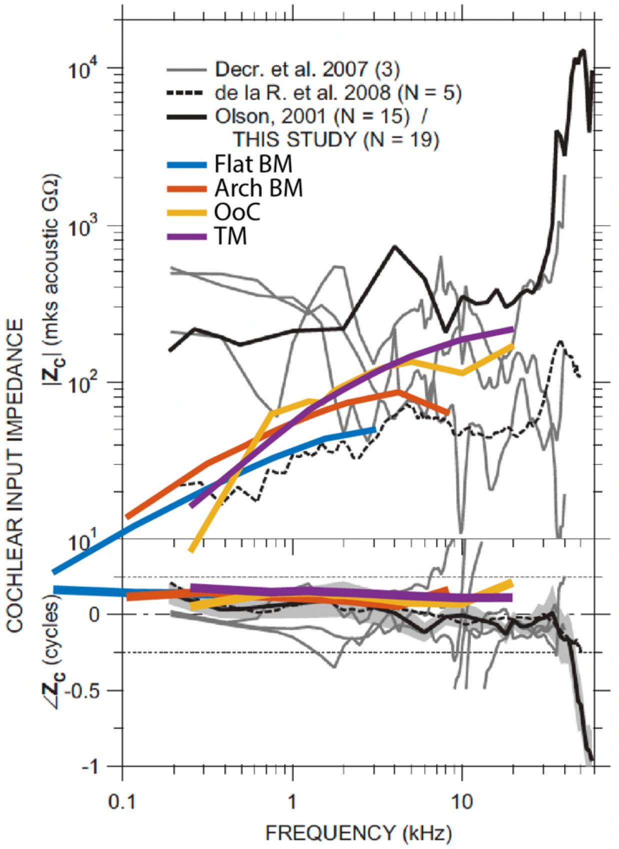


FIG. S5. (Color Online). Cochlear input impedance $Z_C$ of the Mongolian gerbil compared to full-length model morphological variations. Magnitude (upper) in acoustic GOhms ($10^9$ Pa s/m$^3$) and phase (lower) in cycles are plotted in grayscale from 100 Hz to 60 kHz ('THIS STUDY' in the legend refers to the origin of this figure (Ravicz et al., 2008)). Results from the current investigation are overlaid in color for each of the four variations considered: flat BM (blue), arch BM (orange), pseudo-OoC (yellow), and TM (purple). (Modified from (Ravicz et al., 2008)).

The experimental data displayed varied magnitudes with some animals showing a largely decreasing or flat frequency response while others increase slightly from 100 Hz to 30 kHz and rise dramatically above 30 kHz. The absolute value also differed between studies covering roughly one order of magnitude across frequencies. However, this variation could be an underestimate as the data plotted from de la R. (De La Rochefoucauld et al., 2008) was a mean of five animals under different experimental conditions. The conditions most similar to normal hearing with a sealed bulla cavity and covered pars flacida showed a similar frequency dependence to the mean but a magnitude lower by a factor of four (not shown). The phase was near 0.1 cycles up to 5 kHz, near -0.1 from 5 to 30 kHz, and decreased sharply above 30 kHz reaching -1 cycle by 60 kHz. However, Ravicz et al. described the phase as roughly zero below 30 kHz consistent with a resistance-dominated impedance.

The cochlear input impedance of each model variation in our investigation showed a similar frequency dependent magnitude which increased by around a factor of ten over roughly two orders of magnitude in frequency compared to a factor of three for the averages of de la R. and Ravicz (i.e. Olson/This Study). The absolute magnitude largely overlapped experimental results, spanning approximately 10 to 100 acoustic GΩ ($10^9$ Pa s/m$^3$). The phase was around 0.15 cycles across frequencies for all cases, similar to experimental results

With each added morphological feature, the impedance at the highest frequencies tested increased. The high frequency impedance has been attributed to the square root of the partition stiffness at the base (for review, see Puria and Allen, 1991). From the flat BM to the TM/ISS model, $Z_C$ increases approximately by a factor of five suggesting a stiffness increase near a factor of 25. Using our static studies of the middle turn as a proxy, we found the stiffness increased by a factor of 38 across the same two models. It should be noted that this impedance was based on theoretical calculations which did not include the sharp increase for basal most frequencies as measured in gerbil above 30 kHz. However, we also did not explore model responses above this frequency. At the lowest frequencies, the input impedance is influenced by the stiffness of the round window membrane. Nedzelnitsky (1980) reported this took effect below 300 Hz with Lynch et al. (1982) claiming the round window impedance dominates below 40 Hz in the cat. The model did not include a round window membrane and instead enforced a zero-pressure boundary condition on the round window area. However, using round window stiffness estimates from cat and a stiffness-dominated impedance inversely proportional to frequency, the round window impedance would intercept the plot at 16 GΩ and 0.16 kHz—hardly overlapping with responses from any model variation.

**E. Calibrating cochlear responses relative to ear canal pressure**

The current model does not explicitly include a middle ear which would have allowed us to calculate OoC motions for a specific ear canal pressure. Instead, the local OoC displacements and cochlear pressure gains were determined empirically using multiple transfer functions that incorporate the oval window pressure, stapes velocity, and ear canal pressure. The local transverse BM displacement $x_{BM}$ and the local pressure difference about the cochlear partition $\Delta p_{CP}$ are calibrated in the same manner, so we define a new variable $q$ which represents displacement or pressure:

$$q \in \{x_{BM},\ \Delta p_{CP}\}\,. \tag{21}$$

Each full-length model variation was driven by a constant stapes velocity generating the stapes-normalized response $q/v_{ST}$. Multiplying by the middle ear admittance $Y_{ME} = v_{ST}/p_{EC}$ then gives the gain relative to ear canal pressure $q/p_{EC}$ so

$$\frac{q}{v_{ST}} \cdot \frac{v_{ST}}{p_{EC}} = \frac{q}{p_{EC}}\,. \tag{22}$$

The displacement gain $x_{BM}/p_{EC}$ and pressure gain $\Delta p_{CP}/p_{EC}$ frequency responses shown in panels E and F of Figs. 4-7 were calculated using Eq. 22.

Using the specific cochlear input impedance $z_C = p_{OW}/v_{ST} = A_{fp}Z_C$ with $A_{fp}$ the stapes footplate area, we could further subdivide the stapes-normalized responses:

$$\frac{q}{p_{OW}} \cdot \frac{p_{OW}}{v_{ST}} = \frac{q}{v_{ST}}\,. \tag{23}$$

Combining Eq. 22 and 23 gives

$$\frac{q}{p_{OW}} \cdot \frac{p_{OW}}{v_{ST}} \cdot \frac{v_{ST}}{p_{EC}} = \frac{q}{p_{EC}}\,. \tag{24}$$

The full-length model displacement and pressure difference responses, cochlear input impedances, and middle ear transfer function (based on Ravicz et al. (2008)) are shown in Fig. S6. The main text Fig. 9D & 9E presents the displacement and pressure difference relative to ear canal pressure up to a factor of 1.3 above the BFs of each model.

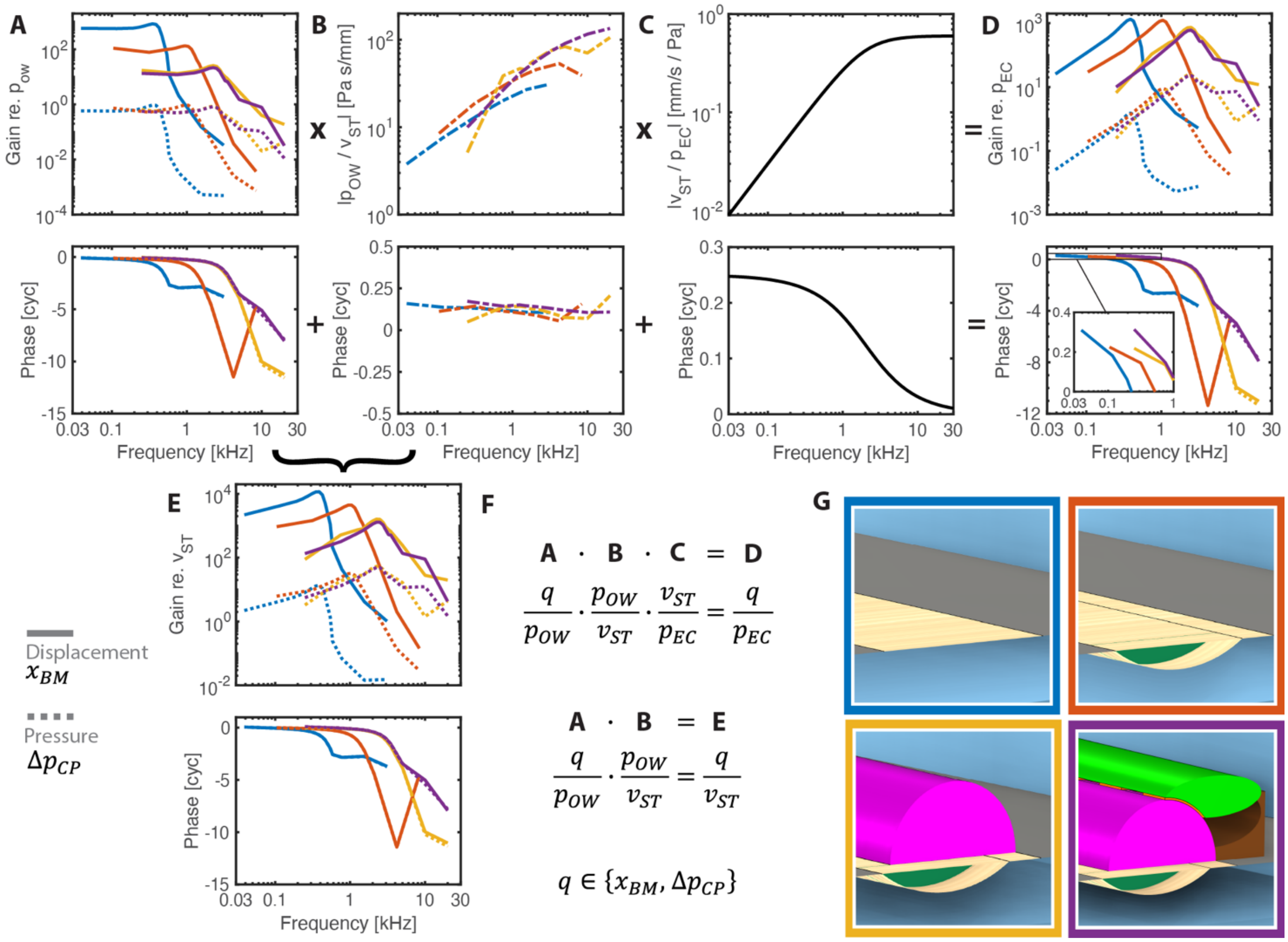


FIG. S6. (Color Online). Calculation of transverse BM displacement and local cochlear partition pressure gain frequency responses. Results are colored for each of the four partition variations considered: flat BM (blue), arch BM (orange), pseudo-OoC (yellow), and TM (purple). Displacement measurements are shown as solid lines and pressure measurements as dotted lines. Displacement phase unwrapping was known from the spatial response. By definition, a positive pressure difference produced a negative transverse displacement, so displacement phase was shifted by 0.5 cycles in all plots to offset this definition. Pressure difference phase was unwrapped to within +/- 0.5 cycles of the displacement phase for comparison. A-E) Magnitude (top) and phase (bottom) of each transfer function: A) local displacement, pressure normalized to oval window pressure $q/p_{OW}$ (nm/Pa and unitless, respectively); B) specific cochlear input impedance $z_C = p_{OW}/v_{ST}$; C) middle ear admittance $Y_{ME} = v_{ST}/p_{EC}$; D) displacement, pressure normalized to ear canal pressure $q/p_{EC}$ (nm/Pa and unitless, respectively); E) displacement, pressure normalized to stapes velocity $q/v_{ST}$ (nm s/mm and Pa s/mm, respectively). The cochlear input impedance (B) was calculated for each variation and used for both displacement and pressure gain from a given variation as indicated by the four colored, dot-dash lines. F) Equations corresponding to the top row of plots (A-D, Eq. 24) and the left half of plots (A, B, E; Eq 23). G) Partition morphologies: flat BM (top left, blue border), arch BM (top right, orange border), pseudo-OoC (bottom left, yellow border), and TM (bottom right, purple border).

The displacement and pressure frequency responses overlapped in magnitude and phase across variations for most frequencies. Relative to stapes velocity (bottom left panels), the responses were bandpass. When subdivided using the oval window pressure, part of the response frequency dependence can be seen to come from the input impedance. Relative to oval window pressure, the responses were lowpass-like with a slight increase near BF for each respective variation. The pressure gain was close to one for low frequencies before decreasing rapidly at frequencies beyond the BF. This was indicative of pressure from the oval window propagating apically until the BF region, regardless of what frequencies that corresponded to.

The middle ear admittance $Y_{ME}$ had a low frequency magnitude slope of 6 dB/octave and phase of one-quarter cycle before approaching 0.6 mm/s/Pa with a 2 kHz cutoff frequency. Together with the displacement and pressure responses and input impedances, this resulted in a gain re. ear canal pressure for any given variation whose low frequency slope was nearly proportional to frequency squared and whose phase was one-quarter cycle. From the flat BM to the arch BM variation, the pressure increased and displacement remained constant re. BF. On the other hand, the pressure increased slightly and displacement decreased slightly re. BF between the arch BM and OoC variations. While these first appear as different reactions to a change in effective stiffness, they are well accounted for considering the responses relative to oval window pressure and the cochlear input impedance in addition to the middle ear admittance.

#### F. Calibration of a Passive Slice Model

Slice models, lacking a stapes input, necessitate distinct stimulation methods compared to full-length models. Nevertheless, frequency responses must be calibrated for comparison to data obtained experimentally. Wang and Puria (2025) evaluated slice model responses by comparing them to a corresponding full-length model, so the driving differential pressure was known. Detailed slice models do not have corresponding full-length models and cannot take advantage of this approach but perhaps can be approximated. The transfer functions presented in the preceding section offer useful insight for this problem.

The middle ear admittance and cochlear input impedance have been studied for common laboratory animals and humans. The differential pressure in a passive cochlea relative to oval window pressure (Fig. S6A dotted lines) is also well described as the input pressure propagates apically until the best frequency region, regardless of the absolute frequency. Therefore, one can reconstruct the passive differential pressure for an arbitrary ear canal input pressure by working in reverse to Eq. 24[3]. By calibrating the slice model stimulation to produce this local pressure difference, one can drive the slice model in a similar fashion as a local section of a full-length model.

The primary challenge to implementing this approach is the determination of "best frequency" for a slice model since it is required for the differential pressure transfer function. The most reliable method we found for this task could be divided into three steps: 1) calculate the "resonant frequency" for a flat BM version of the cochlear slice model using equations for a plate bending in fluid, 2) determine the wavenumber at this frequency through eigenfrequency analysis of the flat BM version, and 3) calculate the BF for the detailed cochlear slice model using this wavenumber and an eigenfrequency analysis of the detailed slice model. We found this to work in part because the wavenumber relative to BF was largely invariant between morphological variations (main text Fig. 9H), meaning we could use the wavenumber at BF from a flat BM slice model to estimate the BF for a detailed slice model. While not trivial, the BF and wavenumber at BF could be more readily estimated for a flat BM slice model.

The flat BM model constitutes a plate bending in fluid. Puria and Steele (2008) presented an equation for the resonant frequency of this system which we found was 1.3 times higher than the measured BF for the flat BM model. This was near a half octave higher and therefore approximated the in vivo, active BF for the slice model. Specifically, the formulation equated the impedance of a hinged plate under transverse deflection with the impedance of an inviscid, incompressible fluid, which could be rewritten using the plate stiffness as

$$f = \frac{1}{\pi}\sqrt{\frac{S_{\mathrm{Vol}}}{\pi\rho}} \tag{25}$$

where $S_{\mathrm{Vol}}$ is the stiffness (applied BM pressure over displaced transverse area) and $\rho$ is density assuming equal plate and fluid density. Although the solution for the clamped condition could not be as readily calculated[4], the solution for hinged conditions is considered reasonable based on results from the four partition variations, described below.

Three-dimensional fluid coupling is required to properly calculate the dispersion relation for a flat BM cochlear box model (main text Fig. 4G) and this coupling was properly captured using an eigenfrequency analysis of the Floquet slice model (Fig. 4I). Particularly considering anatomically accurate scalae of unique shapes and asymmetries, performing an eigenfrequency analysis of a flat BM version of one's detailed slice model is the most reliable way to calculate the model's dispersion. From this wavenumber-frequency relationship, one can pair a single wavenumber (first bending mode) with the resonant frequency from Eq. 25.

Considering the wavenumber relative to BF as largely invariant between morphological variations, as seen Fig. 9H, one can complete the final step to estimate the BF for an arbitrary cochlear slice model. Performing an eigenfrequency analysis on the slice model with all its detail will produce multiple propagating modes (Fig. 8). However, one will correspond to the classical traveling wave mode. By taking the wavenumber corresponding to the flat BM resonant frequency and the new traveling wave dispersion curve, one can then estimate the in vivo BF for the slice model.

Results were consistent when the method was applied to the model variations explored in this study. The flat BM variation stiffness gave a resonant frequency 1.3 times higher than the measured BF for the flat BM study corresponding to a wavenumber of 1.84 cycles/mm. Using the Floquet-derived dispersion curves from the other model variations, we estimated the active BFs for the arch BM, OoC, and TM models to be 1.5, 1.4, and 1.4 times higher than the measured passive BF for each, respectively. This is in very good agreement with the factor of 1.4 (square root of two) roughly expected from passive to active tuning.

#### G. Three-Dimensional Fluid Coupling

There is an ongoing debate on the necessity of considering fluid coupling in three dimensions and even whether responses with three dimensional (3D) fluid coupling vary substantially to those with two dimensional (2D) fluid coupling (Steele and Taber, 1979; Kolston, 2000; Lim and Steele, 2002; Zweig, 2015; Altoè and Shera, 2020; Altoè, 2026). Taking our flat BM variation—most similar to the analytical cases of the past—we found the predicted 2D dispersion relations failed to capture the wavenumber as measured from the phase of the full-length 3D model (Fig. 3F&G). However, we also found that when including the mass of the partition tissues and enclosed fluids, the 2D approximation was sufficient for predicting the dispersion of the variations with additional morphological features. Here, we first compare the dispersion relations predicted using several 3D cochlear hydrodynamics studies (Steele and Taber, 1979; De Boer, 1981; Elliott et al., 2011) against the dispersion from our 3D full-length flat BM model. Then, we explore the 3D approximation when applied to the other partition variations with tissues surrounding the BM.

##### *1. Comparing 3D Approximations*

When the BM moves, an adjacent layer of fluid moves with it. The equivalent height of this layer—$h_{eq}$ (also referred to as $Q$)—adds an effective mass to the partition and depends on the local wavelength of the traveling wave and the dimensionality of fluid coupling. Although our flat BM variation included length-dependent parameters like scala height and BM width, the analyses of $h_{eq}$ discussed below were constructed for uniform box models.[5]

Steel and Taber (1979) explored 3D fluid coupling using the Wentzel-Kramers-Brillouin (WKB) approximation[6] in comparison to experimental data from physical models with length-dependent parameters. The equivalent height for a BM clamped on both ends (radial profile proportional to $\sin^2(y)$) with equal, square cross-section scalae was[7]

$$h_{eq,3D} = \frac{2B}{3}\left(\frac{1}{kH\tanh(kH)} + 2\sum_{j=1}^{\infty}\frac{(H/j\pi B)^2[\sin(j\pi(B+C)/H)+\sin(j\pi C/H)]^2}{[1-(jB/2H)^2]^2 m_j H\tanh(m_j H)}\right) \tag{26}$$

where

$$m_j^2 = \left(\frac{j\pi}{H}\right)^2 + k^2 \ . \tag{27}$$

Here, $B$ is BM width, $k$ is wavenumber, $H$ is scala height (equal to scala width), and $C$ is distance from scala wall to BM edge. Summation was terminated when the maximum difference between two calculations was less that 1%[8]. In the 2D approximation where the wavelength is long compared to the scala width, the first term in Eq. 26 dominates the sum leaving $h_{eq,2D} = \frac{2B}{3}\left(\frac{1}{kH\tanh(kH)}\right)$. In the 1D approximation where the wavelength is also long compared to the height, the equation reduces further to $h_{eq,1D} = \frac{2B}{3}\left(\frac{1}{k^2H^2}\right)$.

To determine the dispersion relation, the authors equated the time-averaged kinetic and potential energies such that

$$\omega = \sqrt{\frac{4S_{\text{Vol}}\Gamma}{2\rho h_{eq}+\rho t_{bm}}} \ . \tag{28}$$

where $S_{\text{Vol}}$ is stiffness and $\Gamma$ relates to the potential energy of the plate. The denominator sums fluid effective mass and (negligible) plate mass where $\rho$ is density (equal for plate and fluid) and $t_{bm}$ is the BM fiber thickness. Stiffness was calculated by evaluating the slice model deformation in response to static pressure, as shown in Fig. S3. For comparison, this was within 10% of the of the value computed for an orthotropic plate with clamped edges as presented by the authors: $S_{\text{Vol}} = K = 8\pi^4 E t_{bm}/B^5$ with $E$ the transverse Young's modulus. Defining orthotropy as the ratio of the elastic modulus in the longitudinal direction to the modulus in the transverse direction $\sigma = E_l/E_t$, one can determine $\Gamma$ for an orthotropic plate with clamped edges. Multiplying $\sigma$ by the wavenumber-dependent terms in $\Gamma$ for an isotropic plate with clamped edges gives

$$\Gamma = \frac{B}{6}\left\{1 + \sigma\left[\frac{1}{2}\left(\frac{kB}{\pi}\right)^2 + \frac{3}{16}\left(\frac{kB}{\pi}\right)^4\right]\right\} \ . \tag{29}$$

The dispersion relations using the 1-, 2-, and 3D equivalent height approximations and Eq. 28 are shown in Fig. S6A.

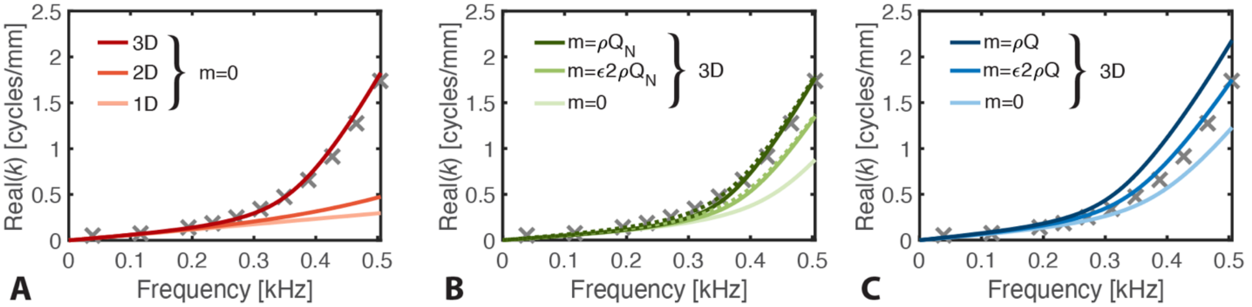


FIG. S6. (Color Online). Comparing three approaches to computing dispersion relations with three-dimensional fluid coupling. The dispersion of the full-length, flat BM variation (inviscid) is shown with x-marks at each frequency the model was solved (up to a factor of 1.3 times the BF of the viscous model). Left) Dispersion based on (Steele and Taber, 1979) obtained using 1-, 2-, and 3D approximations for equivalent height in light to dark lines, respectively. The 3D approach closely matched the data with no additional mass loading. B) Dispersion curves based on (Elliott et al., 2011) consider 3D fluid coupling with variable added mass (in addition to the effective fluid mass inherent to 3D coupling). The fractional width of the BM is $\epsilon = B/W$. Dotted lines indicate the dispersion of mass calculated using the entire equivalent height $Q$ instead of just the near field height $Q_N$, providing a more direct comparison with (C). Close overlap was found when half the mass of the fluid was added ($\rho Q$ or $\rho Q_N$). C) Similar to the center plot, each dispersion curve, based on (De Boer, 1981), considers 3D fluid coupling with variable added mass. De Boer's formulation did not separate near and far field components, so only added mass proportional to $Q$ was calculated. Using this formulation, the tapered, full-length, flat BM model dispersion (x-marks) was most similar to the 3D dispersion with an added a mass of $\epsilon 2\rho Q$.

The 1D approximation (Fig. S6A, light line) is linear in frequency with a slope that matches the low frequency dispersion relation calculated from the full-length, flat BM variation. While the 2D approximation does increase the wavenumber at higher frequencies, the difference pales in comparison to the difference changing from two to three dimensions. The 3D approximation (Fig. S6A, dark line), which considers the partition stiffness and fluid loading, is nearly identical to the dispersion found from the full-length model (x-marks).

Elliot et al. (2011) explored fluid coupling in a discrete cochlear model based on the work of Steele and Taber. Following the derivation for clamped edge conditions in (Elliott et al., 2010), an identical equivalent height can be derived as in Eq. 26, which we will now call $Q$. The fluid coupling impedance can be expressed directly proportional to the equivalent height with

$$Z_{FC}(k) = 2i\omega\rho Q(k) \ . \tag{30}$$

The authors then decomposed the fluid impedance and thus $Q(k)$ into near and far field components. The far-field component satisfied the 1D, long-wave approximation with

$$Q_F = \frac{1}{k^2 h} \tag{31}$$

where $h = 3H^2/2B$ for clamped boundary conditions. The near field $Q_N$ was defined to include all terms of the coupling impedance aside from the far-field term, so $Q_N = Q - Q_F$.

The dispersion relation was obtained by equating the fluid and partition impedance. From (De La Rochefoucauld and Olson, 2007) we get

$$Z = -i\omega\left({}^{S}/_{\omega^2} - m_{solid} + {}^{Ri}/_{\omega}\right). \tag{32}$$

Combining Eq. 30 and Eq. 32 with damping $R$ set to zero gives

$$2i\omega\rho Q(k) = -\left(i\omega m_{solid} + {}^{S}/_{i\omega}\right) \tag{33}$$

with damping set to zero. Here, $m_{solid}$ is the mass of the tissues on the partition and $S = S_{\text{Vol}}B$. Solving for frequency gives the dispersion relation

$$\omega = \sqrt{\frac{S}{2\rho Q(k) + m_{solid}}} \tag{34}$$

with curves presented in Fig. S6B.

The calculated dispersion with no added mass ($m_{solid} = 0$) did not overlap that of the full-length model. This is because the dispersion relation from Eq. 34 differed from that of Eq. 28 by $\sqrt{4\Gamma/B}$, which turns out to be a necessary factor in computing the 3D dispersion for an orthotropic plate (both solutions were for clamped edges). Interestingly, the effects of orthotropy and plate bending could be compensated for by instead adding mass to the partition—setting $m_{solid} \neq 0$. The added mass required to replicate the data was 50% of the original fluid mass due to coupling (i.e. $m_{solid} = \rho Q$). No substantial difference was found between adding fluid mass from the near field ($Q_N$) versus the entire field ($Q$) (solid versus dotted lines, respectively).

Finally, we considered the three-dimensional solution of De Boer (1981). Taking the fractional width of the BM to be $\epsilon = B/W$, the equivalent height for hinged edge conditions was defined

$$Q = \frac{\epsilon}{k\tanh(kH)} + \frac{\pi}{2}\sum_{j=1}^{\infty} \frac{4\epsilon\cos(j\pi\epsilon)}{\pi(1-4j^2\epsilon^2)} \frac{\sin(j\pi\epsilon)}{j\pi\epsilon} \frac{1}{p_j\tanh(p_j H)} \tag{35}$$

where

$$p_j^2 = \left(\frac{2j\pi}{H}\right)^2 + k^2 \ . \tag{36}$$

The results from using the dispersion relation in Eq. 34 with $Q(k)$ replaced by Eq. 35 are shown in Fig. S6C. We found the dispersion with no solid mass again did not match that measured from the full-length model. However, it was closer compared to the previous approach which used same dispersion equation with equivalent height from Steele and Taber (Fig. S6B, $m = 0$). Since the baseline result was closer, less added mass was required to replicate the full-length model dispersion. We found the response was best matched by scaling the fluid mass by the fractional width of the BM ($m = \epsilon 2\rho Q$).

#### 2. *3D Approximations of Partition Variations*

Puria and Steele (2008) argued that given appropriate treatment of 3D fluid coupling, experimental results could be well described without considering the mass of the tissues attached to the BM. In the previous section, we found the dispersion predicted using the WKB approximation from Steele and Taber (1979) for an orthotropic, clamped plate with no added mass was nearly identical to the dispersion found in our tapered, full-length, flat BM variation. Since we had three other full-length models with cochlear partition variations including tissue surrounding the BM, we tested their hypothesis that dispersion depends on stiffness alone and there is no significant mass attached to the BM.

Since the model parameters were unchanged between partition variations (e.g. scala height, BM width, etc.), the only modification to the dispersion calculation was to update the effective stiffness of the partition calculated using the deflection in response to static pressure as seen in Fig. S3. The results are shown in Fig. S8. The response to the flat BM variation (Fig. S6A) is reproduced in Fig. S8A along with the dispersion from the viscous flat BM model (filled circles). For the arch BM variation, the 2D approximation overlapped the measured dispersion (Fig. S8B and main text Fig. 5G). This was despite the BM width occupying the same fraction of the scala width as in the flat BM model—slightly greater than 1/4. The 3D approximation in this case greatly overestimated the wavenumber for a given frequency—about a factor of five higher than the measured wavenumber near BF. Considering the OoC partition variation, the measured dispersion of the full-length model fell in between the calculated 2D and 3D fluid coupling approximations (Fig. S8C). At BF, the wavenumber was roughly 1.5 times higher than the wavenumber predicted using the 2D approximation and 2.3 times lower that the 3D approximation. Similar to the flat BM partition variation, we found the wavenumber-frequency relation of the TM variation overlapped well with the 3D fluid coupling approximation (Fig. S8D). Across all variations, the predicted low frequency, long-wave wavenumber matched with the measured wavenumbers. The primary differences were in the short-wave region at frequencies leading up to and beyond BF, roughly starting one to a half-octave below BF.

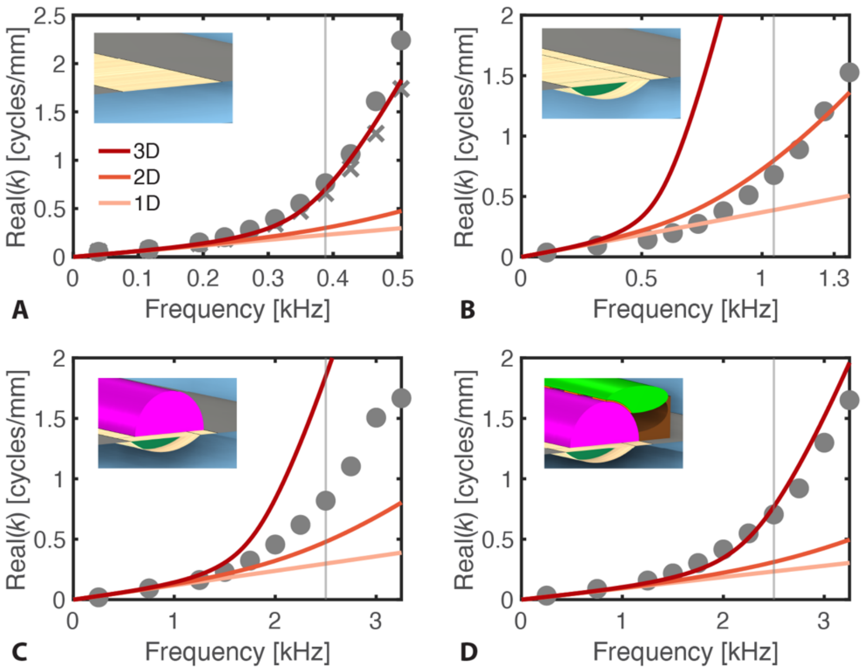


FIG. S8. (Color online). Three-dimensional fluid coupling approximations based on (Steele and Taber, 1979) compared to dispersion measured across the four partition variations: flat BM, arch BM, OoC, and TM. 1-, 2-, and 3D approximations are indicated by the light to dark lines, respectively. Dispersion measured from the full-length models are indicated by markers—circles for models with viscosity and crosses for inviscid models (flat BM only). Data are plotted up to 1.3 times the BF of the viscous models with BF indicated by tick marks. Insets in the top left of each plot show the partition morphology corresponding to the data in that plot. A) Flat BM. The 3D wavenumber approximation for an orthotropic, clamped plate in inviscid fluid was nearly identical to that measured in the full-length model. B) Arch BM. The measured dispersion was best predicted by the 2D approximation. C) OoC. The measured wavenumber increased more rapidly in the short-wave region than the 2D approximation predicted, but not as rapidly as the 3D approximation. D) TM. The 3D fluid coupling approximation best matched the dispersion measured from the full-length model with the most partition morphological features explored in this study.

Here we compared the dispersion from 1D to 3D longitudinal fluid-coupling approximations of a uniform box model with an orthotropic, clamped plate in inviscid fluid considering solely the plate stiffness to the dispersion of tapered box models with varying degrees of morphological detail surrounding the cochlear partition in viscous fluid. We found the 3D approximation was remarkably close to the dispersion from the most simple and most detailed partition morphologies we explored (flat BM and TM variations, respectively). However, we also found the 3D approximation did not predict the dispersion of the two intermediate morphological variations (arch BM and OoC). Importantly, the approximation overestimated the wavenumber in both cases. The mismatch could be corrected by considering a difference between the calculated and effective stiffness, damping, tissue-based longitudinal coupling, or additional mechanisms although more work is required to determine the proper accounting. Since mass is theorized to increase the wavenumber for a given frequency beyond the effects of stiffness alone, this suggests that the mass of tissues surrounding the partition is negligible. While we found the mass of the tissues and enclosed fluids predicted the dispersion across variations using a 2D fluid-coupling approximation, those effects could be partly explained by using 3D longitudinal fluid coupling.

### FOOTNOTES

[1] The phase decrease was faster than quadratic, as is assumed in the short-wave region considering only partition stiffness. See section IV.G. for a detailed description of the 3D dispersion.
[2] Higher order bending profiles were seen at frequencies four and eight times higher than the BF of a given variation (not shown). All portions of this manuscript and supplement concern passive mechanics, so it remains to be seen whether active amplification applied to any partition variation would excite these or other bending modes, as has been suggested (Lin et al., 2024).
[3] Inherent to this approach is the assumption of a particular middle ear and cochlear impedance. As seen in Fig. S6, the cochlear input impedance can be well approximated as a resistor with no frequency dependence and the middle ear admittance as a high pass filter. This simplifies the application and removes nuances from experimental conditions.
[4] $\text{Sin}^2(x)$ is not an eigenfunction of the second- or fourth-derivative operators unlike sin(x), so equating the plate and fluid impedances retains dependence on radial position along the plate.
[5] Length-dependent parameters break the spatial translation invariance of the uniform box model, which introduces nonlocal coupling. There do not exist explicit, scalar $h_{eq}(k)$ for nonuniform models, hence the WKB and numerical approximations.
[6] Also known as the Liouville-Green approximation. See Frost (2024) for a comprehensive review.
[7] The equation was written in (Steele and Taber, 1979) with $\sin(j\pi/L_1)(b+c)$ which should be $\sin(j\pi(b+c)/L_1)$.
[8] Since the numerator of even order terms was effectively zero, iteration differences were calculated between every other index. The threshold was reached after including 11 terms in the sum.
